# GABOLA: A Reliable Gap-Filling Strategy for *de novo* Chromosome-Level Assembly

**DOI:** 10.1101/2021.09.07.459217

**Authors:** Wei-Hsuan Chuang, Hsueh-Chien Cheng, Yu-Jung Chang, Pao-Yin Fu, Yi-Chen Huang, Ping-Heng Hsieha, Shu-Hwa Chen, Chung-Yen Lina, Jan-Ming Ho

## Abstract

We propose a novel method, GABOLA, which utilizes long-range genomic information provided by accurate linked short reads jointly with long reads to improve the integrity and resolution of whole genome assemblies especially in complex genetic regions. We validated GABOLA on human and Japanese eel genomes. On the two human samples, we filled in more bases spanning 23.3Mbp and 46.2Mbp than Supernova assembler, covering over 3,200 functional genes which includes 8,500 exons and 15,000 transcripts. Among them, multiple genes related to various types of cancer were identified. Moreover, we discovered additional 11,031,487 base pairs of repeat sequences and 218 exclusive repeat patterns, some of which are known to be linked to several disorders such as neuron degenerative diseases. As for the eel genome, we successfully raised the genetic benchmarking score to 94.6% while adding 24.7 million base pairs. These results manifest the capability of GABOLA in the optimization of whole genome assembly and the potential in precise disease diagnosis and high-quality non-model organism breeding.

Availability: The docker image and source code of GABOLA assembler are available at https://hub.docker.com/r/lsbnb/gabola and https://github.com/lsbnb/gabola respectively.

## Introduction

Since the advent of genomic sequencing technology, for 20 years, the attempt to resolve the human genome has continued by improving techniques. In 2003, the completion of the human reference genome was marked as a triumph in biomedical research^1^. The advances of sequencing techniques, from the initial Sanger Sequencing to next-generation sequencing (NGS) allowing generation of high throughput sequencing data, and toward the third-generation sequencing platform characterized by long read sequences, offers great possibilities to accomplish accurate human genomes^2^. The latest human genome version, GRCh38, was released in 2013. GRCh38 reference genome has hitherto represented the basis for all human genomic studies that identify functional genic regions, define regulatory elements, compare genomic diversity, conduct populational genomics analyses, and search for disease-causing mutations^3^. In addition, as the cost of sequencing drops, millions of people have already been sequenced and their DNA sequences differed against the human reference genome to identify genetic factors associated with health and disease. The current practice in genomic analyses is to align the reads generated by the sequencers to GRCh38 reference genome to determine the location of the reads, and in turn, construct the individual’s genome. Nevertheless, GRCh38 reference genome is still susceptible to undesirable problems which haven’t been fixed.

### Characteristics and Current Problems in Human Reference Genome

The latest genome build GRCh38 remains several problems in application and is still incomplete with numerous unsolved gaps. First, GRCh38 is an underrepresented genome for entire human populations. GRCh38 is a composite genome originated from limited donors with certain genetic ancestry, Caucasian and African especially^4^. Therefore, genetic diversity existing in other populations is not captured, in that the resolution to detect variants would be restrained. Although the supplement of alternate contigs have been provided through different versions to increase the diversity, studies still find limitations when using GRCh38 as a reference in populations other than European and North American populations. Specifically, some regions of the human genome consist of structurally complex sequences that are highly variable among populations or even individuals due to diverse ancestry background^5, 6^. The underrepresentation of human genome will cause great burden, so-called reference bias, in alignment-based genome assembly process, since the reads carry alleles other than reference alleles would have lower mappability, thus, higher chance to be falsely assembled.

Second, it has been reported that highly repetitive sequences are universal in human genome^7^. Therefore, megabase-scale long range information is required to provide large structural information in the genome. Next generation sequencing (NGS) technology, which GRCh38 reference genome mainly based on, is performed by random fragmentation of DNA molecules, giving rise to short read sequences (200-300 bp) that are solvable by sequencer^8^. This type of sequencing data is accurate but limited in length, a fatal constraint in solving highly repetitive sequences and large structural variants. Consequently, GRCh38 reference genome inherits this limitation and the ability and accuracy to discern these structural variants are still concerning issues in genomic studies. Especially, a large portion of intergenic regions, known as the dark matter in the genome, consisting of myriad repeats was basically not addressable. Despite the development of Third-Generation Sequencing (TGS), Long reads with read length of over hundreds kilobases may cover genomic information in length but with the cost of notoriously high error rate^9^ . This issue becomes magnificent in disease diagnosis, such as cancer, since error-prone repair processes causing genetic instability are prevalent in cancer genomics. Moreover, the process of error correction of long read sequences is laborious and painstaking. The computational cost is so high that it is still impractical to adopt as a routine pipeline.

Third, GRCh38 is still incomplete with numerous unsolved gaps^10–12^. These gaps are primarily located in the centromeric regions and the short arms of the acrocentric chromosomes. The missing of genome content has been contributed to the existence of near-identical tandem repeats^5^. These regions are reported to be associated with cellular protein production machinery and to be crucial in ribosomal function. Due to the absence of content, a significant portion of whole-genome sequencing (WGS) reads from each individual is discarded because they have low mapping quality when aligned to the human reference genome, even though the discarded reads are not contaminants^13^. Consequently, aligning reads to human reference genome does not provide the true configuration of these regions in the sequenced individual. These collectively cause considerable biases in variant calling based on current human reference genome.

#### Toward Precise Personalized Genome

The alternative of alignment-based genotyping is to de novo assemble individual’s genome. Without accepting the parti pris from reference genome, de novo assembly can benefit the genotyping process in two ways: 1. gaining new sequence assemblies for previously unreported genomic regions and 2. characterizing the individual’s genome in a reference unbiased fashion^14, 15^. As a result, gaps existed in reference genome may be resolved, and individual’s private variants can be identified rather than being discarded. The increment of the content of personal genome would pave the road to identify unforeseen variants and facilitate disease-association test.

Technologies provide long-range genomic information with acceptable accuracy should be incorporated in characterising large structural mutations in disease diagnoses. Although some synthetic long-read techniques using barcodes to provide long-range information, such as 10xGenomic sequencing (10xG), they inherit the similar limitation as NGS in highly repeated genomic region^16^. One possible solution is to incorporate optical mapping (OM) in sequencing pipeline. When integrated with other sequencing reads, OM provides the order and orientation of sequence fragments, identification and correction of potential chimeric joins, and estimation of the gap size between adjacent reads^17^.

#### Supernova 10xG and Bionano

Standard short read sequencing generates accurate base-level sequence to provide short range information, but struggles to contribute to long-range information. Moreover, short reads are inaccessible to the region of high identity repeats and paralogs. Therefore, we adopted 10XG Linked Reads combined with Bionano optical mapping to gain a comprehensive genomic scaffold.

10XG Linked Reads technique utilised molecular barcodes to tag reads and provide long-range genomic information and phasing SNVs, indels, and structural variants^18^. The key concept is that by introducing a unique barcode to every short read deriving from a few individual molecules, thus short reads can be linked together. Linked-Reads provide long range information at length of 50kb, which is a significant improvement compared to Illumina short-read sequencing (range information of 300 bp).

Supernova assembler utilised read pairs to reach over the short gaps. Barcode is informative to fill the large gaps. If physical location of two scaffolds is actually in proximity, then with high probability, multiple molecules in the partitions bridge the gap between two scaffolds. We show that the linked-reads produced with 10X Genomics Chromium chemistry and assembled with Supernova assembler generated scaffolds with an N50 of 45 Mbp with the longest individual scaffold of 129 Mbp (Table 1).

**Table 1.**
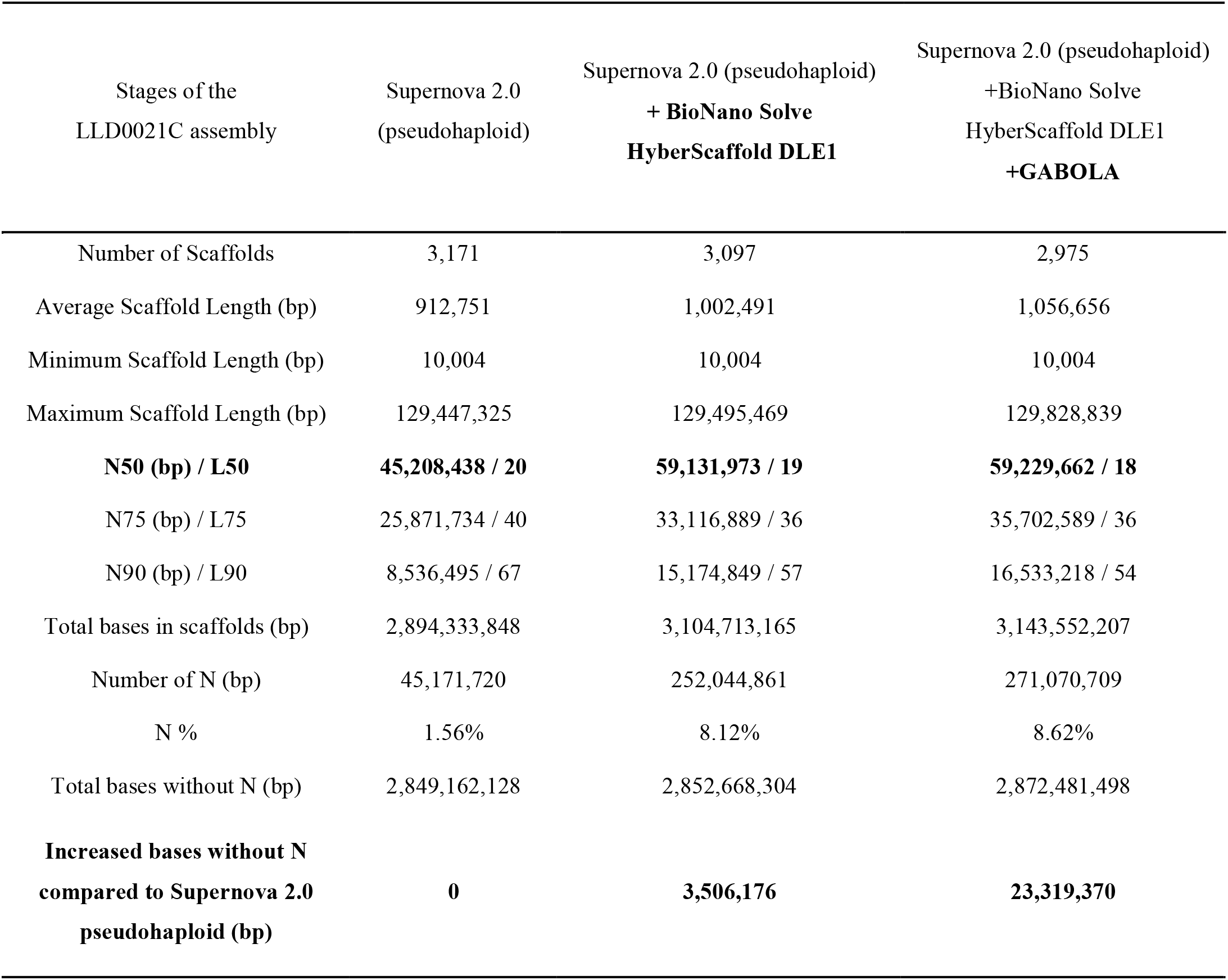
Assembly statistics show notable increase especially in N50 and total base length on sample LLD0021C after GABOLA

| Stages of the<br>LLD0021C assembly | Supernova 2.0<br>(pseudohaploid) | Supernova 2.0 (pseudohaploid)<br>+ BioNano Solve<br>HyberScaffold DLE1 | Supernova 2.0 (pseudohaploid)<br>+BioNano Solve<br>HyberScaffold DLE1<br>+GABOLA |
| --- | --- | --- | --- |
| Number of Scaffolds | 3,171 | 3,097 | 2,975 |
| Average Scaffold Length (bp) | 912,751 | 1,002,491 | 1,056,656 |
| Minimum Scaffold Length (bp) | 10,004 | 10,004 | 10,004 |
| Maximum Scaffold Length (bp) | 129,447,325 | 129,495,469 | 129,828,839 |
| **N50 (bp) / L50** | **45,208,438 / 20** | **59,131,973 / 19** | **59,229,662 / 18** |
| N75 (bp) / L75 | 25,871,734 / 40 | 33,116,889 / 36 | 35,702,589 / 36 |
| N90 (bp) / L90 | 8,536,495 / 67 | 15,174,849 / 57 | 16,533,218 / 54 |
| Total bases in scaffolds (bp) | 2,894,333,848 | 3,104,713,165 | 3,143,552,207 |
| Number of N (bp) | 45,171,720 | 252,044,861 | 271,070,709 |
| N % | 1.56% | 8.12% | 8.62% |
| Total bases without N (bp) | 2,849,162,128 | 2,852,668,304 | 2,872,481,498 |
| **Increased bases without N**<br>**compared to Supernova 2.0**<br>**pseudohaploid (bp)** | **0** | **3,506,176** | **23,319,370** |

To further provide long-range information for disentangling complex genomes, we adopted Bionano optical mapping. Bionano optical mapping set specific 6-mers sequence motif as markers to provide a blueprint of the genome structure^17^. Hybrid scaffolds were generated by merging the bionano optical map and in silico sequence map generated from the Supernova assembly in order to validate the assembly results, order and orient sequence fragments, identify potential chimeric joins, and help estimate the gap sizes between adjacent sequences. When combined with Bionano Genomics optical mapping, the scaffold N50 increased to 59Mbp (Table 1). For a total of 116 hybrid scaffolds, the longest of which was 110 Mbp (Table 1). The complementarity of linked-reads and optical maps is likely to make the production of higher quality genomes more routine and economical, whereas we still find myriad gaps and private insertions that are unresolved.

#### Current Pitfalls in Supernova Assembler

Although de novo Genome Assembly may be free of reference biases, there are several technical issues in assembly problems, including GC content biases and difficult-to-solve repeat regions^19^. The GC content bias originated from sequencing process which would result in uneven sequencing depth across genome. Most of de Bruijn graph-based assemblers use the read depth information for constructing contigs and scaffolds. Thus, the uneven sequencing depth impedes the genome assembly. De Bruijn-based assemblers use an average coverage cutoff threshold for contigs to prune out low coverage regions, which tend to include more errors. (large gap average size) Secondly, as mentioned in the Characteristics and Current Limitation of Human Reference Genome, approximate 50% of the human genome involves nonrandom repeat elements, including long interspersed nuclear elements (LINEs), short interspersed nuclear elements (SINEs), long terminal repeats (LTRs) and simple tandem repeats (STR)^20^. Repeats cause disarrangements or gaps in the assembly, and also cause a nonuniform read depth, thus resulting in copy loss or gain in the assembly^21^.

#### Summary of GABOLA and results

Here, we designed our method, “De Novo Genome Assembly Based On Localization And Local Assembly Of Scaffolds For Precision Health And Genome Breeding (GABOLA)”, based on 10xG barcoded linked-reads and de novo local assembly. It consists of four main modules: 1) Local-Contig-Based (LCB) Gap Filling”, 2) Global-Contig-Based (GCB) Gap Filling , 3) Global-Contig-Based (GCB) Scaffolding and 4) Local-Contig-Based (LCB) Scaffolding. While GCB Gap Filling and LCB Gap Filling fill in the vacant sequence regions to create a more complete genome, GCB Scaffolding and LCB Scaffolding extend scaffold lengths in the hope of reaching the chromosomal level. These four algorithms all share the common core concept described as follows: We first use BWA-mem to map all linked-reads to the draft assembly and select those that are aligned to the flanking of gaps or ends of scaffolds with high quality. Then, we assemble contigs with all reads that belong to the same barcodes as the previously selected reads. In the case of GCB Gap Filling and GCB Scaffolding, we take long reads or scaffolds as input which allows us to skip over the de novo assembling process aforementioned. With these contigs, we identify the best qualified contig or collinear contigs to fill in corresponding gaps or connect scaffolds into longer ones.

We performed GABOLA separately on different species such as the Japanese eel for evaluation and conducted complete experiments on two human samples. The workflow tested on the two human individuals, LLD0021C and CHM13, varied slightly. With the former, we adopted Bionano optical mapping to elongate the scaffolds before running GABOLA on the assembly. Whereas with the latter, no additional assembling techniques other than Supernova was applied. Despite the differences, the results both show significant increase in gene content and growth in sequence length. We discovered up to 23Mb and 46Mb of new content compared to the initial Supernova output respectively in LLD0021C and CHM13, including those non-reference unique insertions located in exonic and regulatory regions. With the help of Bionano optical mapping, we were able to stretch the maximum scaffold length of sample LLD0021C to over 130Mb and the number of N50 from 45,874,826 to 59,222,212 base pairs. We also saw similar improvements in CHM13, our methods successfully extended the maximum scaffold length by 11Mb and N50 by 4.9Mb. Both results indicate that GABOLA can indeed identify personal variations involving structural variants and segmental duplication with minimal gaps, providing better genomic resolution and improving diagnosis based on personal genome with low cost and high accuracy.

## Results

### 1. Testing GABOLA on one Taiwanese human sample

We experimented with different approaches on an individual, LLD0021C, including interchanging GABOLA module orders, applying different filtering criteria and integrating other scaffolding methods. Finally, we reached the ideal version of our assembly following the pipeline depicted below.

#### I. Pipeline description for LLD0021C analysis

**Figure 1.**
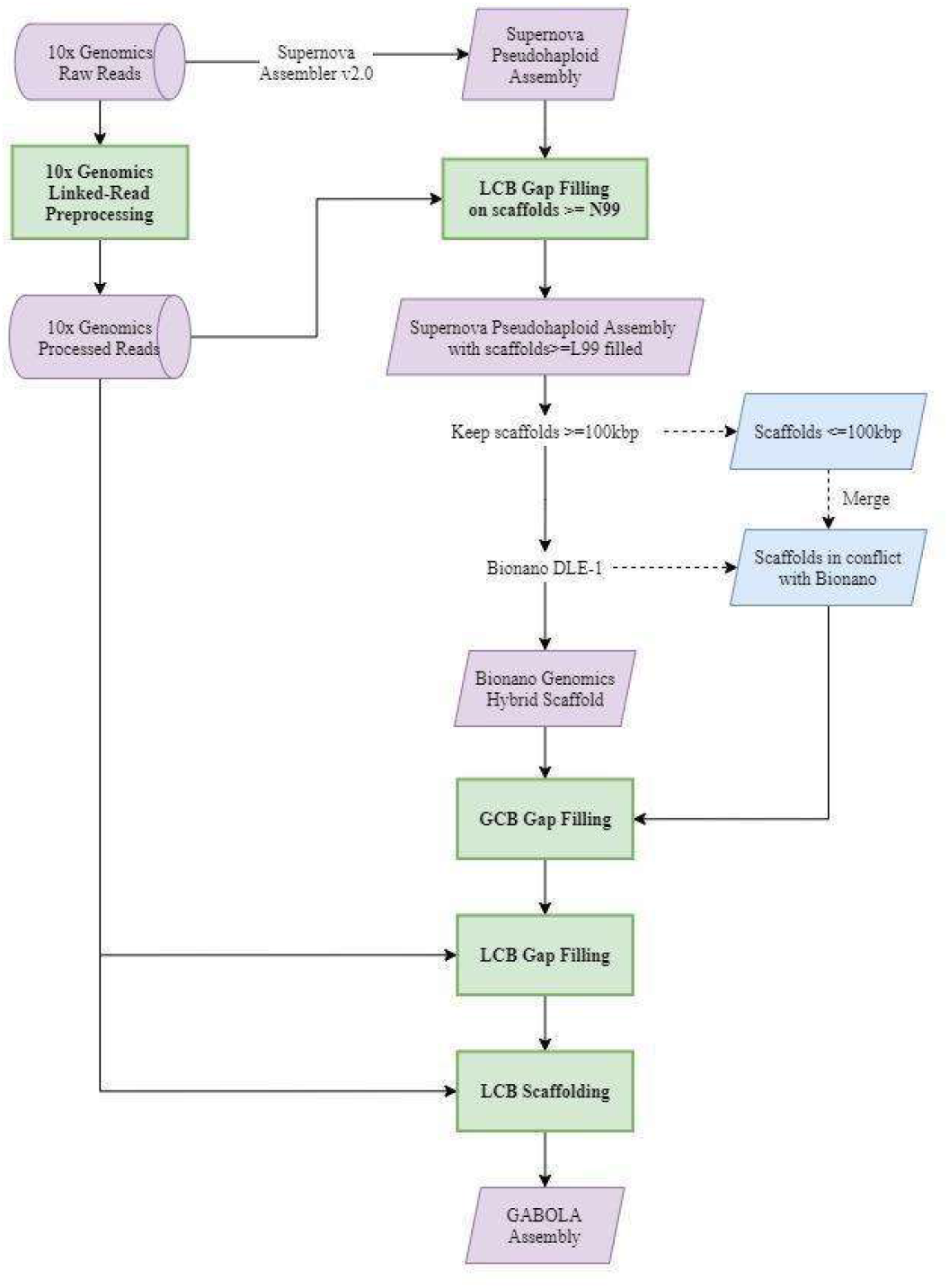
The pipeline of our method for LLD0021C.

The whole experiment done on sample LLD0021C can be divided into two stages: The first one is the incorporation of existing sequence assemblers. The second one is the application of our method, GABOLA.

Preliminarily, we generated a haplotype draft assembly of 3,171 scaffolds by running the 10x Genomics Software Supernova v2.0^19^ using raw linked-reads of the human sample dataset, LLD0021C (Supplementary Note 1). We then incorporated the optical mapping technique of Bionano Genomics^17^ to extend the lengths of our scaffolds (Supplementary Note 2). We decided to fill gaps on a subset of the assembly containing 1,171 scaffolds longer than the N99 length (22,459 bp) before the Bionano DLE1 protocol with LCB Gap Filling. The Bionano process only accepts scaffolds with lengths greater than 100kbp as input; therefore, sequences that are shorter than 100kbp or are in conflict with the Bionano cmap were discarded during the process, leaving 258 scaffolds to undergo the following work. The output comprised a new set of 116 Hybrid Scaffolds.

Subsequently, we entered the second phase of our pipeline by performing GCB Gap Filling, where we managed to fill in larger gaps on the 116 Hybrid Scaffolds with the discarded sequences aforementioned. We retrieved the sequences unused in the previous step and added it back to our whole genome assembly composed of 3,102 scaffolds. Then, we conducted LCB Gap Filling on the assembly, where we filled gaps by the unique set of contigs assembled for each gap with linked-reads. Finally, we connected 237 scaffolds via LCB Scaffolding, yielding a final draft assembly of 2,975 scaffolds.

#### II. Comparing Supernova assembler and Bionano to GABOL

Regarding the Supernova haploid draft and the final draft, we filled in 23,319,370 new base pairs in total (Table 1). Among those gaps, 5,785 were completely-filled and 3,992 were partially-filled. As shown in Supplementary Table 1, we mainly resolved gaps under 1 kbp in both cases. We observed that even though the mean of completely-filled gaps may not be outstanding, the mean of partially-filled gaps (4,274.27 bp) could be improved by raising the number of iterations in the last step of gap filling (see Discussion). In addition, the largest gap size for both types of gaps are 59,608 bp and 100 Kbp, indicating the capability of our method to bridge bigger gaps.

We also raised the number of N50 from 45,208,438 bp to 59,229,662 bp with an increment of 14 Mbp. The maximum scaffold length increased by 381,514 bp to approximately 130 Mbp, which is longer than half of the human chromosomes and akin to the length of chromosome 11 (135,186,938 bp).

#### III. Seeking validation through scaffold-to-reference alignments

For further assessment, we aligned our final assembly to the globally-used reference genome,GRCh38.p13^22^, and Human Diversity Reference (HDR)^23^, via minimap2^24^. The total covered bases on GRCh38.p13 calculated using bamcov (https://github.com/fbreitwieser/bamcov ) is 2,782,616,483 with the percentage of 90.10%; while the covered bases of HDR is 2,787,140,499. This result exhibits a solid relation between our assembly and the references, but the remaining uncovered bases still implies the need for a Taiwanese genome reference.

#### IV. Gain of genome content in GABOLA compared to Supernova assembly

To demonstrate that GABOLA can gain higher resolution in functionally important genomic regions, we found out the sequences that are filled by GABOLA but not in Supernova assembly (GABOLA-filled sequence). GABOLA-filled sequences are then annotated based on GENCODE in GRCh38 coordinate^25^. We classified GABOLA- filled sequences into four main GENCODE biotypes: coding sequence, non-coding transcript, pseudogene, and others. (Table 2). We discovered that GABOLA retained sequences in over 28 thousand protein coding regions, meaning that these sequences with direct functional impact have been ignored in previous assembly. Specifically, GABOLA significantly improved genome contents in genes located in pericentromeric and telomeric regions, including sequences in exons and transcripts (Table 3). In non-coding transcript, lncRNAs are long regarded as important regulatory element for gene expression. In GABOLA, 3,552 additional sequences in lncRNA were identified. Moreover, sequences in rRNA (rDNA array) are the most difficult-to-solve regions in genome assembly problems. We found that GABOLA has the ability to resolve sequences encoding rRNA. Throughout evolutionary process, pseudogenes are generated through genome duplication and retrotransposition. Therefore, pseudogenes are composed of duplicated and repeated sequences that may be difficult to assemble. We found GABOLA improved the genome content in approximately three thousand pseudogenes. In general, GABOLA outperformed in multiple functional classes and these newly assembled genome contents would significantly affect the prediction of protein coding and regulation.

**Table 2.** Filled sequence in GABOLA compared to Supernova assembly categorized by GENCODE biotype

| **Biotype** | **Counts of Filled Genomic Regions** |
| --- | --- |
| **Coding Sequence:** | 28,861 |
| Protein coding | 28,746 |
| Immunoglobulin and T cell receptor | 115 |
| **Non-coding Transcript:** | 3,768 |
| Long non-coding RNA (lncRNA) | 3,552 |
| Non-coding RNA (ncRNA) | 206 |
| tRNA | 7 |
| rRNA | 3 |
| **Pseudogene:** | 2,996 |
| Unprocessed pseudogene | 2,328 |
| Processed pseudogene | 504 |
| Unitary pseudogene | 78 |
| rRNA pseudogene | 9 |
| Other pseudogene | 77 |
| **Others** | 44 |
| To be experimentally confirmed (TEC) | 44 |

**Table 3.**
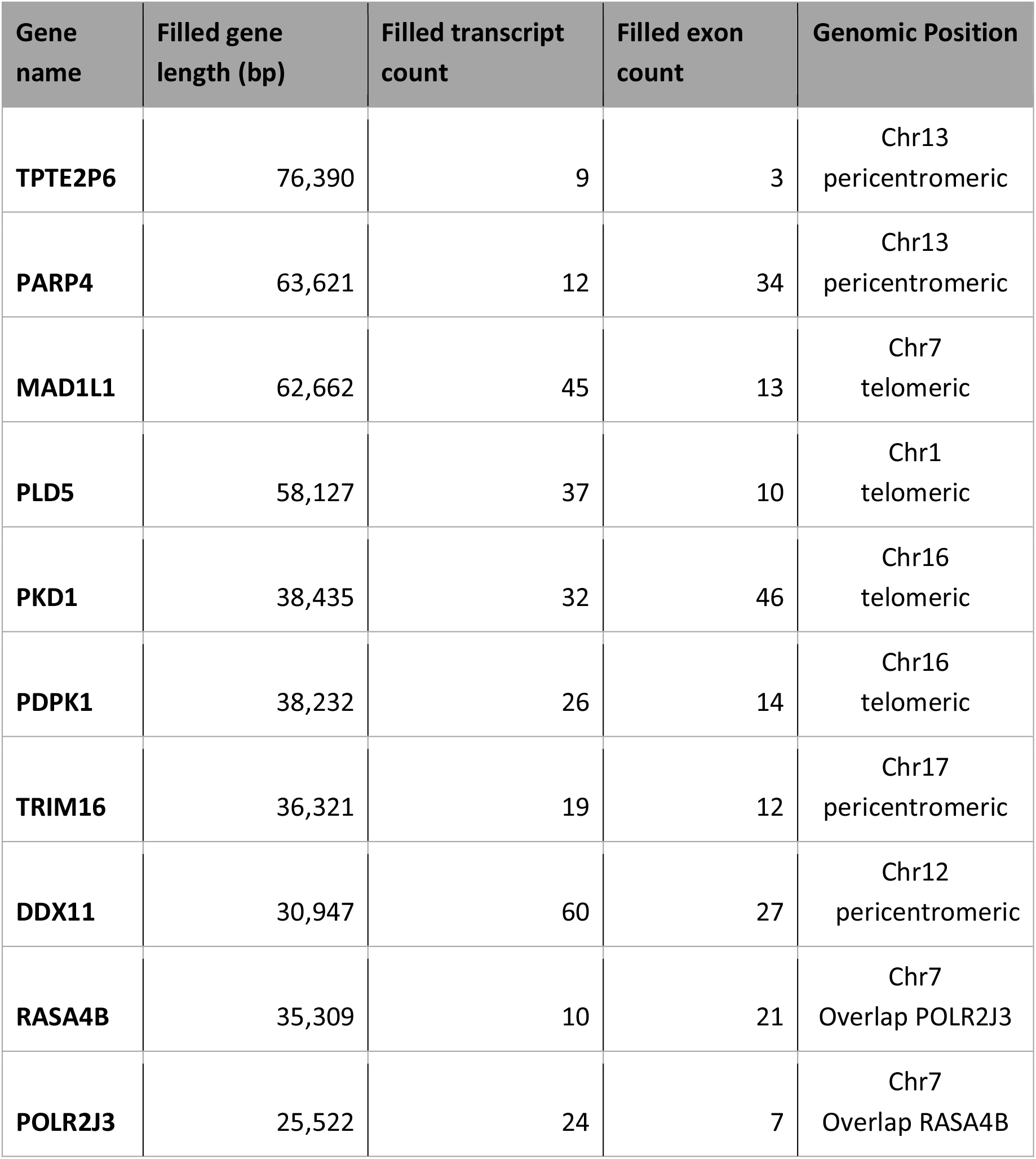
GABOLA significantly improves genome contents in protein coding genes located in pericentromeric, telomeric regions, and difficult-to-solve regions.

#### V. The performance of Supernova and GABOLA in masked reference region (N in reference genome)

There are still genomic regions marked as gap of “N” in GRCh38 Human reference genome, especially in homologous centromeric and genomic repeat arrays. These regions with repeated sequence are notoriously known for bad resolution in read-alignment based and short-read de novo assembly. We find that GABOLA is able to extend the scaffolds to fill genomic contents in gap regions, accounting for 462,705 bp in 12,046 gap regions , and performs significantly better than Supernova assembly.

#### VI. GABOLA outperforms Supernova Assembly in Repeat Regions

Assembling contigs and scaffolds in the highly repeat regions has been the error-prone in de novo genome assembly. However, repeat elements account for a considerable percentage (approximately 45%) of homo sapien genome. Therefore, we assessed the performance of GABOLA in these repeat regions. We use RepeatMasker^26^, which is based on Repbase, to identify the repeat regions. When compared to Supernova assembly, 11,031,487 additional bases were masked and identified as repeat elements. 1,138,402 bp and 5,626 bp were classified into SINE and LINE categories, respectively. (Table 4) Surprisingly, 218 repeat patterns were identified in GABOLA assembly but not in Supernova.

**Table 4.** GABOLA outperformed supernova in repeat sequences

| Region | Supernova Length | GABOLA Length | Increased Percentage |
| --- | --- | --- | --- |
| **SINEs:** | 360,307,214 | 367,599,283 | 2.02% |
| ALUs | 297,675,946 | 304,339,130 | 2.24% |
| MIRs | 62,062,069 | 62,685,814 | 1.01% |
| **LINEs:** | 555,422,256 | 570,384,310 | 2.69% |
| LINE1 | 467,877,986 | 481,834,046 | 2.98% |
| LINE2 | 78,456,751 | 79,370,127 | 1.16% |
| L3/CR1 | 7,146,074 | 7,219,478 | 1.03% |
| **LTR elements:** | 235,284,651 | 241,576,774 | 2.67% |
| ERVL | 48,026,801 | 49,125,199 | 2.29% |
| ERVL-MaLRs | 97,217,164 | 99,198,073 | 2.04% |
| ERV_classI | 78,440,079 | 80,985,565 | 3.25% |
| ERV_classII | 8,176,299 | 8,798,235 | 7.61% |
| **DNA elements:** | 86,856,844 | 88,266,581 | 1.62% |
| hAT-Charlie | 36,794,256 | 37,280,601 | 1.32% |
| TcMar-Tigger | 32,699,038 | 33,364,831 | 2.04% |
| **Unclassified:** | 4,164,825 | 4,329,007 | 3.94% |
| Small RNA | 1,171,956 | 1,235,812 | 5.45% |
| Satellites | 9,122,975 | 14,774,390 | 61.95% |
| Simple repeats | 36,143,467 | 39,011,022 | 7.93% |
| Low complexity | 6,216,364 | 6,361,304 | 2.33% |

#### VII. Evaluation of GABOLA’s scaffolding methods

We sought to verify the accuracy of scaffolding, to find out whether the candidate scaffold pairs we determined based on barcode positioning agree with the standard references. Therefore, we did an examination on the set of scaffolds connected by GABOLA. First, we extracted those joined by scaffolding, namely “New Scaffolds (NS)”, and the scaffolds composing the NS set, which will be termed “Original Scaffolds (OS)”. We saw that 237 OS were concatenated into 115 NS after the Scaffolding process: 105 NS consisted of 2 OS and the other 10 consisted of 3 OS. Next, we aligned all NS to GRCh38.p13 with minimap2. 4 NS were unmappable, which could be caused by the disparity between individuals, faults produced during sequencing, the error occurred when joining scaffolds or them possibly being new found sequences. After filtering out supplementary hits and keeping the one with the highest Mapping Identity, we got the best alignments of each NS. This gave us the position information we needed to cut out the regions of reference sequences with bedtools getfasta^27^. We then mapped the OS to the reference sequence fragments clipped according to their respective NS, for we believed that the OS should align perfectly to the regions if they were supposed to be connected. As the result manifests, the 8 OS from the 4 unmappable NS were also unable to adhere to GRCh38.p13, eliminating the possibility of Scaffolding being the reason for this finding. As for the other scaffolds, 182 OS aligned to their corresponding segments, thus proposing that 87 NS are connected correctly. We went on to scrutinize the remaining 24 NS and filed them under two occasions: The first is the case where the scaffold pair selection was indeed correct but were joined in the wrong orientation during Scaffolding, this could most likely be triggered by some algorithmic defects we failed to notice. Fortunately, only 7 NS fell under this category. The second is the case of wrong scaffold pair selection, possibly arising from false barcode selection or genetic differences between people. There were 17 NS paired incorrectly, which is tolerable since 87 were correct. We did a double-check on the 4 unmapped NS and 24 possibly wrong NS on HDR to validate this outcome. Surprisingly, the results are entirely the same as GRCh38.p13.

To further study these unmapped NS, we applied Augustus^28^ to make gene prediction followed by protein BLAST^29^ to investigate if the predicted sequences are conserved in organisms. Consequently, we found 11 inferred genes in 4 NS. Among these genes, 2 genes are found to be homologs of the existed genes, PTZ00395 and DUX4. Specifically, DUX4 is located within a repeat array in the sub-telomeric region of chromosome 4q, where the similar repeat array is also found on chromosome 10.

Taken together, the results still suggest that our induction of candidate scaffold pairs based on barcode positioning and execution of scaffold connection via Scaffolding is in fact reliable.

### 2. Validating GABOLA through CHM13 htert cell line

With the previous sample, LLD0021C, we took various measures to optimize the assembly. While the results show extreme potential, we sought to demonstrate the capability of GABOLA alone. From the preceding experiment, we inferred a pipeline which we believe would be exemplary for generating a high-quality assembly. We then evaluated it through the genome assembly originated from the CHM13 htert cell line.

#### I. Description of the CHM13 assembly pipeline

**Figure 2.**
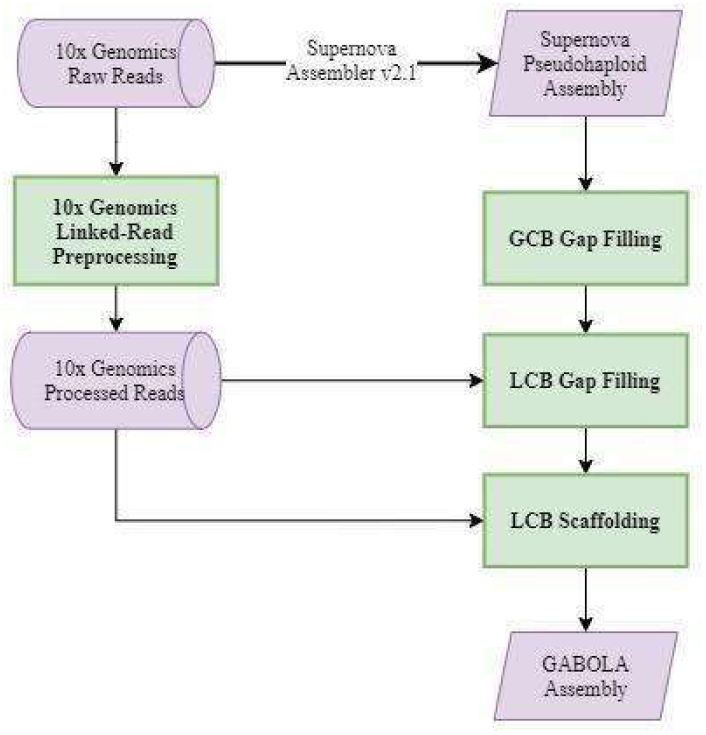
The pipeline of GABOLA for sample CHM13.

Similar to the final workflow done on LLD0021C, we primarily generated a haplotype draft assembly of 4,999 scaffolds by running the 10x Genomics Software Supernova v2.1^21^ using Linked-Reads (Supplementary Note 3). However, the purpose of this experiment is to validate GABOLA, so we did not incorporate Bionano Optical Mapping into this pipeline. We followed the same routine of the second phase of LLD0021C by beginning with GCB Gap Filling, where bigger gaps were mended using the scaffolds assembled by Supernova v2.1. Secondly, we performed LCB Gap Filling on the assembly. Finally, we connected 212 scaffolds via LCB Scaffolding, leaving us with a final draft assembly of 4,787 scaffolds.

#### II. Comparison between Supernova assembler and GABOLA on CHM13 dataset

We discovered 46,287,195 more base pairs in addition to the initial haplotype and managed to reduce the percentage of gaps from 1.21% to 0.87% at the same time (Table 5). We also saw a 11.5Mbp growth in the maximal scaffold length and nearly 5Mbp growth in the quantity of N50. Under further examination, we managed to fill in 18,636 out of 23,349 gaps: 10,768 were completely-filled and 7,868 were partially-filled (Supplementary Table 2). Similar to the results of LLD0021C, the gaps we filled are principally under 1kbp in both circumstances but the maximum size of completely-filled gaps has upscaled to 100kbp. To sum up, even without integrating Bionano we could still gain fruitful results and enhance the assembly to a certain quality.

**Table 5.** GABOLA effectively expands genome size and decreases gaps on CHM13 assembly

| Stages of the<br>CHM13 assembly | Supernova 2.1 (pseudohaploid) | Supernova 2.1 (pseudohaploid)<br>+**GABOLA** |
| --- | --- | --- |
| **Number of Scaffolds** | **4,999** | **4,809** |
| Average Scaffold Length (bp) | 583,041 | 613,731 |
| Minimum Scaffold Length (bp) | 10,000 | 10,000 |
| **Maximum Scaffold Length (bp)** | **91,225,906** | **102,752,858** |
| N50 (bp) / L50 | 39,045,223 / 25 | 44,037,625 / 23 |
| N75 (bp) / L75 | 18,387,696 / 51 | 19,878,974 / 48 |
| N90 (bp) / L90 | 1,578,658 / 108 | 1,799,962 / 101 |
| Total bases in scaffolds (bp) | 2,914,622,463 | 2,951,434,376 |
| **Number of N (bp)** | **35,124,040** | **25,648,758** |
| **N %** | **1.21%** | **0.87%** |
| Total bases without N (bp) | 2,879,498,423 | 2,925,785,618 |
| **Increased bases without N**<br>**compared to Supernova 2.1**<br>**pseudohaploid (bp)** | **0** | **46,287,195** |

#### III. Seeking validation through scaffold-to-reference alignments

We mapped our final draft with minimap2 to the reference genome CHM13v1.0. CHM13v1.0 is the near complete sequence of a human genome constructed by the Telomere-to-Telomere (T2T) consortium^30^, it includes over 100 Mbp novel sequences compared to GRCh38.p13 with only five known gaps remaining within the rDNA arrays. 91.01% of our sequences were successfully aligned to CHM13v1.0. By closer inspection, we observed that the percentages of coverage were only low on a few chromosomes, including those containing the five unresolved gaps, namely chr15, 21 and 22. This implies that our assembly is pretty close to the well-polished reference, only hindered by the innate flaw of unresolved gaps.

We evaluate the results of GABOLA with the same annotation and repeat elements identification pipeline mentioned above. Likewise, GABOLA has higher performance in all functional classes and repeat elements than Supernova, where GABOLA increases 9.1% genomic content compared to Supernova in CHM13. Notably, GABOLA has a 37.73% increase in exonic regions in CHM13 and 6.69% increase in ncRNA regions. These newly identified genome contents would significantly affect the prediction of protein coding and regulation. The CHM13 genome has inherited characteristics of haploid pattern and higher proportion of disease-related mutations. We find that GABOLA has considerable improvement in CHM13. This may be attributed to the simplicity of the haploid genome and great performance in assembling mutated sequences.

### 3. Performing GABOLA on different species along with Third-Generation Sequencing Techniques

In addition to Next Generation Sequencing, GABOLA is also compatible with Third Generation Sequencing reads. More specifically, an alternative input to de novo assemblies for GCB Gap Filling are long reads. To testify this, we performed our method on one of our genome projects, Anguilla japonica. The draft assembly was based on a design with multiple short read libraries to provide read pairs with ranging information, including pair-end, long- range pairend (1k), and mate-pair jump (2k, 4k, 6k, 8k, 12k and above) libraries using TruSeq/Illumina HiSeq platform. It was first generated with ALLPATHS-LG^31^, SSPACE^32^ and GapCloser^33, 34^. Then, we performed SSPACE and GapCloser with newly supplied PacBio long read contigs generated by Canu^35^. The assembly then underwent multiple other processes, including SALSA^36^ , SSPACE-long^37^ and ALLMAPS^38^. The refined draft assembly contains 4,617 scaffolds with N50 contiguity length 42.6Mbp and total length 1,083 Mbp. We went on to polish this assembly with our own method GABOLA. With PacBio reads and scaffolds assembled from 10x reads by Supernova as input, we conducted three iterations of GCB Gap Filling, each time filtering out used contigs from the previous round. Table 6 depicts a steady growth in the number of N50 and maximum scaffold length. We managed to reduce the percentage of gaps from 4% to 1.76% and increase the gene content by 58M bases in our first try. In terms of quantity, we can say with confidence that the performance of GABOLA is stable and distinctive.

**Table 6.** GABOLA is able to reduce a considerable amount of gaps through several iterations

| Stages of eel genome assembly | original | GCB Gap Filling<br>1st iteration | GCB Gap Filling<br>2nd iteration | GCB Gap Filling<br>3rd iteration |
| --- | --- | --- | --- | --- |
| Number of Scaffolds | 4,617 | 4,617 | 4,617 | 4,617 |
| Average Scaffold Length (bp) | 234,776 | 241,403 | 243,673 | 248,489 |
| Minimum Scaffold Length (bp) | 889 | 889 | 889 | 889 |
| Maximum Scaffold Length (bp) | 73,940,153 | 74,964,264 | 75,377,128 | 76,365,690 |
| N50 (bp) / L50 | 42,610,991 / 10 | 43,649,922 / 10 | 44,030,935 / 10 | 44,552,354 / 10 |
| N75 (bp) / L75 | 25,879,373 / 17 | 25,279,375 / 18 | 25,629,905 / 18 | 26,379,933 / 18 |
| N99 (bp) / L99 | 9,268 / 889 | 10,134 / 881 | 10,368 / 872 | 10,781 / 869 |
| Total bases in scaffolds (bp) | 1,083,962,265 | 1,114,555,594 | 1,125,040,273 | 1,147,271,934 |
| **Total number of N** | **47,921,917** | **19,671,117** | **15,111,710** | **12,579,893** |
| **Reduced number of N** | **0** | **28,250,800** | **4,559,407** | **2,531,817** |
| **N %** | **4%** | **1.76%** | **1.34%** | **1.10%** |
| Total bases without N (bp) | 1,036,040,348 | 1,094,884,477 | 1,109,928,563 | 1,134,692,041 |
| Increased bases without N (bp) | 0 | 58,844,129 | 15,044,086 | 24,763,478 |

We then ran BUSCO v3.0.2^39^ on both the initial and final assemblies for quality assessment. BUSCO evaluates the quality of genome assembly with a predefined gene models of single-copy orthologs which selects orthologous groups of genes present in at least 90% of the species including arthropods, vertebrates, metazoans, fungi, eukaryotes, and bacteria. The completeness of a genome can be estimated by searching those genes in the genome draft which are scanned by using tblastx^40^, hidden Markov models, and Augustus^41^. The predicted genes are marked as “complete” or “duplicate” if the gene models are satisfied in the BUSCO algorithm. Some matches of gene models are partial and are marked as “fragmented”; the remaining BUSCO groups with no matches are marked as “missing”. We use BUSCO to evaluate the genomic contents in our assembly and see if there are more genes being found after running our gap-filling pipeline that those BUSCO groups marked as “fragmented” and “missing” are expected to be recovered after gap-filling.

Using the BUSCO database of taxon Vertebrata, 93.2% of the teleost single ortholog genes (2,409 out of 2,586 genes) are identified as complete genes according to Table 7. Despite the already high score, after three iterations of GCB Gap Filling, we still raised the percentage from 93.2% to 93.7%. To further enhance our assembly, we introduced a polishing tool aimed at error correction, POLCA^42^ . With its aid, we were able to obtain an even higher BUSCO score of 94.6% for this eel assembly.

**Table 7.** GABOLA enhances the quality of eel genome indicated by BUSCOv3.0.2 results

| Stages of eel genome assembly | original |  | 3 iterations of GCB<br>Gap Filling |  | 3 iterations of GCB Gap Filling<br>+ Hi-C data &<br>Polishing Approach |  |
| --- | --- | --- | --- | --- | --- | --- |
|  | % | n | % | n | % | n |
| Complete BUSCOs (C), C=S+D | **93.2%** | 2,409 | **93.7%** | 2,422 | **94.6%** | 2,448 |
| Complete and single-copy BUSCOs (S) | 82.5% | 2,133 | 82.8% | 2,141 | 81.8% | 2,116 |
| Complete and duplicated BUSCOs (D) | 10.7% | 276 | 10.9% | 281 | 12.8% | 332 |
| Fragmented BUSCOs (F) | 2.9% | 76 | 2.4% | 62 | 1.9% | 49 |
| Missing BUSCOs (M) | 3.9% | 101 | 3.9% | 102 | 3.5% | 89 |
| Total BUSCO groups searched | 100% | 2,586 | 100% | 2,586 | 100% | 2,586 |

**Table 8.**
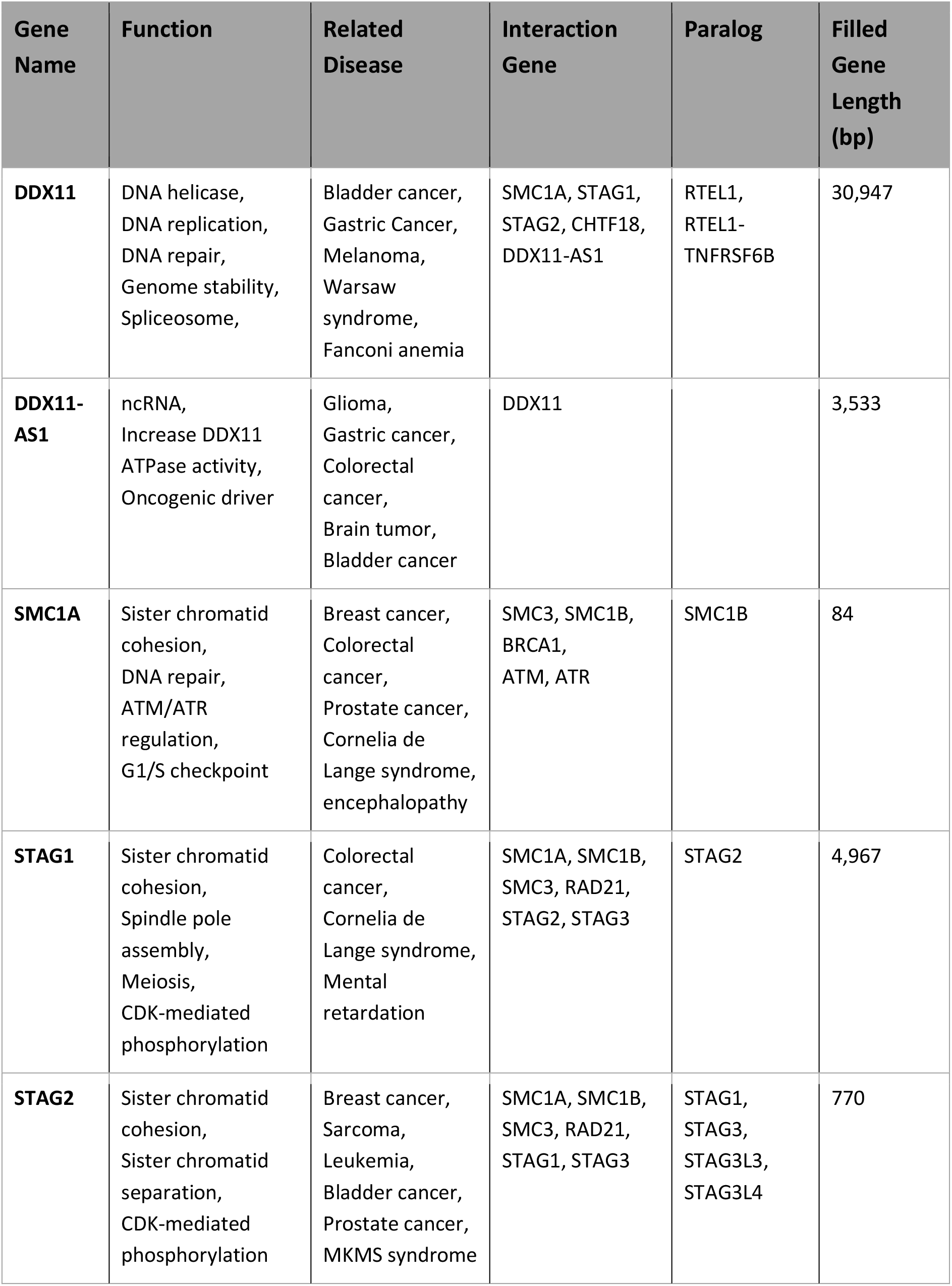
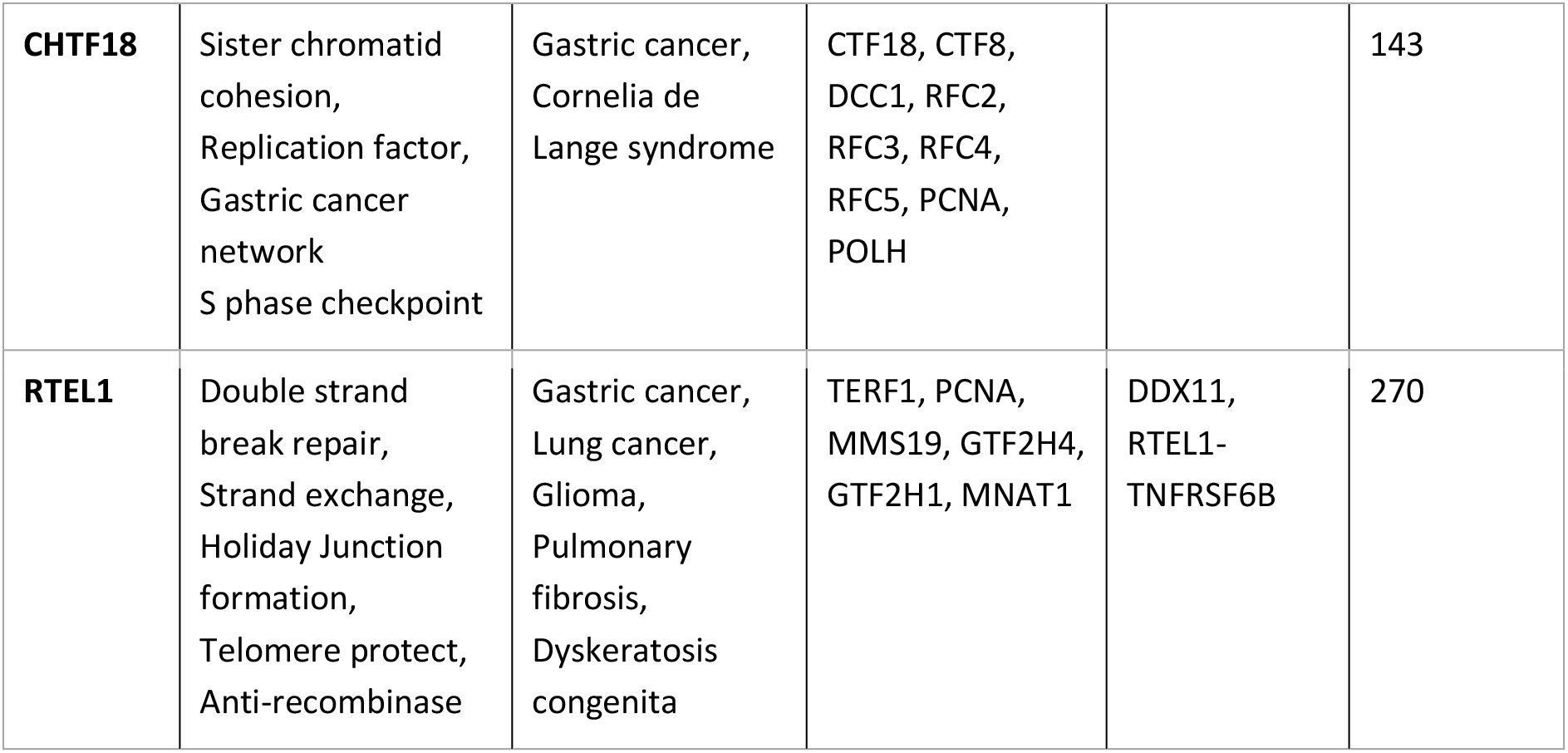
GABOLA is capable of resolving genes related to biological pathways involving complex diseases

## Discussion

We are proposing a de novo genome assembly pipeline, to improve the practice in precision medicine and precision agriculture. GABOLA shows its robustness on combining the advantages from leading sequencing platforms to elongate the resolved sequence while increasing accuracy. GABOLA generates an individual’s complete genome, resolves large structural variants and fills the gap in the current human reference genome. In the validation of GABOLA assembly performance, we focused on genome contiguity and increment of genomic content. In human samples, GABOLA increases N50 from 45Mb to 59Mb, and meanwhile, lowers LG75 from 40 scaffolds to 36 scaffolds. We observed an increment of genome contents of over 23 MB in GABOLA when compared to Supernova assembly. These increased genomic content strongly improve the resolution in functionally important regions and duplicated sequences, as we found abundant GABOLA-filled sequences in protein coding regions, long non-coding RNA and pseudogenes. GABOLA, moreover, identified over 0.4 Mb novel sequences in gap regions in the GRCh38 human reference genome. It is worthwhile to note that GABOLA assembly features its ability to relocate the reads in repetitive genomic regions. When comparing the results of repeat sequences, we found additional 11,031,487 bp repeat sequences and surprisingly, 218 repeat patterns based on Repbase database^41^ were identified in GABOLA assembly but not in Supernova. GABOLA also showed its superb performance on other organisms such as Japanese eels, which would powerfully promote genomic studies in non-model organisms.

### Contribution to Human Reference Genome

The standard reference genome is a representative genome that incorporates more or less 10 individuals^43^. For years, a myriad of genomic analyses have been conducted by mapping to this genome, including functional annotation^44^, disease association^45–48^ and genetic ancestry tests^49–52^. However, the nature of mapping restricts its resolution to identify novel variants other than those found in the initially recruited samples^53, 54^. Our proposed GABOLA assembly integrates the merits of linked read technology and optical mapping, while providing our novelty in locating associated reads in potentially difficult-to-solve regions. We showed that despite the usage of short reads, we are able to reconstruct long-range information to the largest extent. Note that our GABOLA assembly is devoid of using the information from reference information, therefore, free from inheriting reference biases.

We discovered GABOLA assembly has significantly improved in filling the gap and increasing gene content in genes related to complex diseases and those involved in important biological pathways, including functions of DNA repair, DNA replication, cell cycle checkpoints, cell signalling transduction, telomeric regulation, etc. These genes are highly related to cell over-proliferation and tumor development, which may give us explicit interpretation in disease mechanisms. Specifically, we discovered GABOLA filled 31 kb additional bases in DDX11 gene compared to Supernova. This gene involves in DNA replication^55^, DNA repair^56^, heterochromatin organization^57^ and cell cyle regulation, interacting with numorous genes related to meiosis and cell cycle checkpoints^58^, including STAG1^59^, STAG2^60^, SMC1A^61^, CHTF18^62^, DDX11-AS1^63^. Theses clusters of genes have been studied to be direct disease-causing factors of several cancers, including breast cancer, colorectal cancer, prostate cancer, gastric cancer, etc. GABOLA increase the content of these genes in hundreds to thousands basepair. The strength of GABOLA also enable the assembly process to distinguish reads of their paralogs. In DDX11 case, GABOLA has the ablility to increase gene content in its paralogs RTEL1 compared to Supernova. We infer that since reads of DDX11 and its paralogs is largely identical in short genomic range, Supernova may misassemble the read to correct genomic position while GABOLA assemble them accurately. In addition, RTEL1, which is functionally important in telomere regulation in tumour development, locates in telomeric region of chromosome 16, making the assembly problem more challenging^64, 65^.

In addition, the linear structure of reference genome has been challenged that it is incapable to present the diversity of human populations^22^. Although several genome projects proposed augment reference with alternative haplotype^66, 67^, it would introduce inefficiency through sequence duplication between the main linear genome and the alternate haplotypes^68^. Meanwhile, the structure of graph genome has been developing and improving. With the graph genome^68–71^, novel variants identified in our assembly could be included in the presentation of genome at the scale of population or family. Through this pipeline, it would improve the accuracy and efficiency of alignment and variant detection.

### Future works of GABOLA assembler

Despite our effort, there are still a handful of unresolved gaps in our assembly. Multiple factors could give rise to this outcome: either the gaps are too big for the contigs to cross over, or the regions are in genetically complex areas such as tandem repeats, or simply due to the lack of DNA information. We have some ideas that may help us enhance our current methods.

The assembling of contigs takes up the majority of the execution time but only the best-aligned contig from the whole set per gap is used in a single run due to the concern of excessive computational cost in our early developing stage. We are aware that the cost ratio is rather lopsided and the rest of the sequences are discarded after a painstaking process of assembly. As stated in Methods, gaps are classified into three groups: completely-filled, partially-filled and unfilled. In the case of partially-filled gaps, there might be more than one contig that is best-qualified and oftentimes we are left with huge vacancies even after gap filling. We believe that by applying iterations on the final step of gap filling we could shorten the gap gradually and make better use of the contig set. Per gap, we would filter out the used contig, align the subset to the flanking and then fill in one or both sides. This process can be repeated several times until no candidate contigs remain.

Reference-guided Scaffolding is another prospect that we have in mind. Although incomplete, the human reference genome data, GRCh38, has been widely used as a standard answer to the homo sapien genome content. We hope that the reference can provide guidance in connecting scaffolds that our method failed to detect and can also act as a validation to our results. What we have in mind preliminarily is to align our scaffolds to GRCh38, and then we gather information about possible scaffold pairs from the alignment, which we will term as “reference-guided scaffold pairs” (RGSP). We assume that sequences that are aligned with high quality within a reasonable distance of one another are supposed to belong to the same DNA fragment, therefore should be connected. There are two ways to concatenate the RGSP, the first is to use the already-assembled contigs from the LCB Scaffolding process in order to save time from generating a whole new set of sequences. The other is by adopting our algorithm: map the linked-reads to the RGSP and assemble contigs from the selected barcodes then find the best hit to connect the scaffolds.

These methods may help us prolong the scaffold lengths to the chromosomal level and the reference to reach its integrity.

### Future application in functional annotation, personalized medicine and genetic ancestry tracing

We still find the immense needs of functional annotation on structural variants. In our analysis, pervasive repetitive elements are identified in the genome, spanning for approximately 45% of the genome, while their functional importance is still unclear. Specifically, 218 repeat patterns are identified exclusively in our assembly and most of them are simple tandem repeats. It is known that several disorders such as neuron degenerative diseases are strongly linked to the repetitive structure of particular genomic segments^72, 73^. Besides, the features of repetitive elements have been exploited to develop genetic markers^74–76^. In the population sense, patterns of repeat sequences could be a useful link to trace the demographic alterations^77, 78^. With these variants annotated, it could be potential markers or even causal variants for disease discovery or ancestry tracing.

Overall, our pipeline is suggested to be a standard practice in the future genomic study. With the availability of inexpensive and accurate reads generated by short read sequencer, a nearly complete genome containing information of private variants are assembled through GABOLA assembly. The genomic data can further map to graph reference genome and then undergo variants analysis via up-to-date variant callers, such as GATK and DeepVariant. Through this procedure, it provides precision and personalized investigation on not only common variants but those difficult-to-detect variants.

## Methods and Materials

### 1. Genomic DNA sample collection of LLD0021C and DNA processing for 10xGenomics linked read sequencing

The linked-read raw data of the human sample, LLD0021C, is provided by Dr. Nancy Wei from Institute of Biomedical Sciences, Academia Sinica. LLD0021C whole blood was obtained. High-molecular-weight genomic DNA extraction, sample indexing and generation of partition barcoded libraries were performed. 1.2 ng of genomic DNA was used as input to the Chromium system. The 10x Genomics barcoded library was sequenced on the Illumina HiSeq2500, yielding a total of 660 million of the raw reads comprising 57x genome coverage.

### 2. Preprocess for Bionano optical mapping of LLD0021C

LLD0021C whole blood was obtained. High-molecular-weight genomic DNA was extracted for genome mapping. Blood was centrifuged at 2000g for 2 min, plasma was removed and sample was stored at 4 ℃. 2.5 µl of blood was embedded in 100 µl agarose gel plugs to give ∼7 µg DNA per plug, using the Bio-Rad CHEF Mammalian Genomic DNA PlugKit. Plugs were treated with proteinase K overnight at 50 ℃. The plugs were then washed, melted and solubilised with GELase. The purified DNA was subjected to 4 hr or drop dialysis to remove inhibitory substances. DNA concentration was determined using Qubit 2.0 Flurometer, and the quality and length were assessed with pulsed-field gel electrophoresis.

The high-molecular-weight DNA was labelled according to commercial protocols using the Saphyr Reagent Kit. Specifically, 300 ng of purified genomic DNA was labeled through the Direct Label and Stain (DLS) method^17^. After labelling, the backbone of fluorescent-labelled DNA was stained with DNA stain (Bionano).

### 3. Genomic DNA sample collection of CHM13

The linked-read raw data of the other sample, originated from the CHM13htert cell line, were obtained from the github website of the Telomere-to-Telomere (T2T) consortium^30^ in the format of raw FASTQ files. Around 1.2 billion reads and 41x effective coverage of data was generated with a mean molecule length of 130 kbp and an N50 of 864 reads per barcode. The draft assembly which we performed our gap filling methods on were generated with Supernova v2.1^21^ using those linked reads.

### 4. Supernova assembler

10XG Linked Reads technique utilised molecular barcodes to tag reads and provide long-range genomic information and phasing SNVs, indels, and structural variants^18^. The key concept is that by introducing a unique barcode to every short read deriving from a few individual molecules, thus short reads can be linked together. The process starts from massive-throughput automated barcoding and library construction by the Chromium System with the input of high molecular weight (HMW) gDNA. The generation of Gel bead in EMulsion (GEM), which encapsulates each tiny micro- reaction in oil, enables barcoding and DNA amplification to occur in a single partition. Input gDNA is fragmented and distributed across 1M partitions. In each partition, the DNA molecules are labelled with a unique 10X Barcode. Approximately 10 HMW gDNA molecules at length of 50kb in each GEM are mixed with barcoded primers and enzymes and undergo isothermal incubation to generate 10x Barcoded amplicons. Therefore, all fragments from the same GEM share a common 10x Barcode. Barcodes recruit short-reads into paralogous gene loci, so Linked-Reads can align reads correctly using barcoded anchors. Linked-Reads provide long range information at length of 50kb, which is a significant improvement compared to Illumina short-read sequencing (range information of 300 bp).

Supernova first demultiplexed molecules and adopted a de Bruijn graph strategy to produce initial genome graph^21^. To improve the genome graph, Supernova utilised read pairs to reach over the short gaps. Barcode is informative to fill the large gaps. If physical location of two scaffolds is actually in proximity, then with high probability, multiple molecules in the partitions bridge the gap between two scaffolds.

### 5. Bionano optical mapping and merging 10xGenomics datasets into a single assembly

Using the Bionano Saphyr instrument, automated electrophoresis of the labelled DNA generated in the nanochannel array of an Saphyr Chip, followed by automated imaging of the linearised DNA. The DNA backbone and locations of fluorescent labels along each molecule were detected using the Saphyr instrument’s software. The length and set of label locations for each DNA molecule defines an individual single-molecule map. Raw Bionano single-molecule maps were de novo assembled into consensus maps using the Bionano IrysSolve assembly pipeline^17^ with default settings, with noise values calculated from the 10xGenomics Supernova assembly.

Bionano optical mapping and 10xGenomics sequencing data were merged together as follows. Bionano assembled contains and the 10xGenomics assembly were combined using Bionano’s Solve tool by identifying the label sites in the sequence based on the labelling specific recognition sequence motifs. Conflicts between the two are identified and resolved, and hybrid scaffolds are generated where optical maps are used to bridge sequence assemblies and vice versa. The resulting combined dataset generated assembly with a scaffold N50 length of 59 Mb. This process reduces 3,171 scaffolds after Supernova assembly to 116 scaffolds, improving assembly quality and accuracy while reducing computation time.

### 6. 10xGenomics Linked-Read Preprocessing by GABOLA

Our method, GABOLA, is composed of one preprocess module and four main modules: “Local-Contig-Based (LCB) Gap Filling ”, “Global-Contig-Based (GCB) Gap Filling”, “Local-Contig-Based (LCB) Scaffolding” and “Global-Contig-Based (GCB) Scaffolding”. To commence GABOLA, the raw reads generated by the 10x Genomics system must undergo preprocessing.

#### 6.1 Classifying and purifying FASTQs

The principal feature of linked-reads is their potential to provide long range genetic information which has been an issue regarding short read sequencing. To optimize this trait, we believe that reads should be categorized by their barcodes so that we could use all reads to assemble contigs in our further work and save computing time.

To do this, we first extract barcode information from each read and append them to their read names in the fastq files with Longranger^79^. Then, we split these fastq files into R1 and R2 reads (Supplementary Figure 1a).

Redundant bases that were attached to the reads during 10x sequencing should be removed. Therefore, 16bp 10x barcode, 6bp random primer, and 1bp of low accuracy sequence were trimmed from an N-mer oligo in R1 reads, and Illumina adapter contaminants are cut from each read pair with the aid of Trim Galore (F Krueger, 2015).

According to the pilot data, we noticed that non-duplicated read pairs delivered a better gap filling result than duplicated read pairs (Supplementary Note 4). Hence, we mark the shorter or the latter read pairs as a duplicate read pair if their mapping positions are in the same start/end site and the sequence of reads are identical to another read pair’s sequence. Duplicated read pairs will be discarded so that every read pair belonging to the same barcode is unique. Finally, we have a list of barcodes each assigned with trimmed and non duplicate read pairs.

#### 6.2 Aligning and Filtering read pairs

The read pairs were then mapped to the genome scaffold set with BWA in end-to-end mode BWA mem^80^ (Supplementary Figure 1b).

We are only interested in read pairs with high mapping quality, so we filtered out secondary, duplicate, supplementary and chimeric alignments to keep the properly aligned read pairs using using sambamba^81^.

Later, we kept those reads with mapping identity greater than 70 and mapping quality with the equivalent of 60. Mapping identity represents the percentage of exact matches of the alignment; a read with mapping identity greater than 70 means that more than 70% of base pairs in that read are mapped to the genome. Mapping quality values are measured by MAPQ values from BWA-mem mappers which represents the possibility of the wrong mapping being negative-logarithmically.

This filtering step ensures that remaining read pairs are aligned with high quality, which in turn provides convincing mapping evidence for subsequent works. We then select barcodes from this high quality read set which is denoted [ReadHQ] in section 4.

#### 6.3 Barcode distribution across scaffolds from alignment results

This step is necessary when conducting Scaffolding, which is the process of connecting two or more scaffolds by inferring the affinity between them.

Based on the filtered alignment result in section 3.2, we generate a list for each scaffold of the barcodes mapped onto its two ends within the given range (default of 20,000bp). We believe that scaffolds who share the same barcodes may imply proximity in their placement on the genome sequence. Thus, we go on to search for scaffolds that share the same barcodes, which we will refer to as “scaffold pairs” in the following text. The scaffold pairs were then recorded in the descending order according to the amount of their common barcodes.

This process yields a list of all barcodes mapped to each scaffold edge and a list of possible scaffold pairs with their shared barcode count.

### 7. Barcode Positioning

As stated in Introduction, short reads from the same partition of 10x GemCode Technology derive from one or few gene fragments and are assigned to a common barcode. Unmapped reads with the same barcode as mapped reads from [ReadHQ] may give us clues about missing DNA information. Therefore, the selection of barcodes plays a crucial role in our methods.

In terms of “Local-Contig-Based (LCB) Gap Filling”, we focus on intra-scaffold gaps . According to the three selection strategies (Supplementary Note 5), we concluded that it is most cost-efficient to assemble contigs with the barcodes picked out in a gap-based fashion. To do so, we initially tackled the task from a larger scale, then narrowed down to each gap per scaffold. Within the high-quality read set [ReadHQ], we gathered those barcodes with a sufficient number (default of 3) of read pairs aligned to each scaffold to generate scaffold-based barcode lists. For each gap, we collected barcodes possessing sufficient read pairs (default of 2) mapped within the flanking. The flanking size varies depending on the gap length to provide more local information about that particular gap.

When it comes to Scaffolding, we are more invested in the barcodes mapped on sequence ends due to the potential of them implying scaffold relations. As stated above in section 3.3, we tallied and recorded all barcodes of the reads that were mapped onto 20kbp within the head and tail of each scaffold. From this barcode information, we generated a list of scaffold end pairs that have the same barcodes and filtered out end pairs that are from the same scaffold. For each scaffold, we kept its corresponding scaffold ends with the top-two shared barcode counts. It will leave us with a final list of scaffolds that have the higher chances of being concatenated, which we will term “candidate scaffold pairs” in the following section 5.3.

### 8. Assembly

#### 8.1 Local-Contig-Based (LCB) Gap Filling

##### I. Contig Assembling

Following barcode selection in the previous section, we now have a unique barcode list for each gap per scaffold. For each barcode in the list, we gather all reads of those barcodes as a pool. Once a barcode has been chosen into the barcode list, all the reads of this barcode will be recruited to help us fill the gaps no matter whether these reads are in a high-quality mapped set [ReadHQ] or not. We use SPAdes assembler^82^ as we mentioned before to generate contigs which we will coin “L-contigs” as in Local-contigs, for they are produced based on barcode localization. Then, all L-contigs are aligned to the target using BWA-mem, and we fill gaps with high contig-scaffold alignment confidence (Supplementary Figure 5a).

##### II. Defining aligned segments of contig-to-scaffold alignments

When we align assembled contigs to the target scaffold, a single contig usually produces more than one alignment. The best alignment is expected as a contig that can fully cover a gap with two high-score aligned segments to the gap flankings. A true alignment in such a case, however, may be interrupted more than once if the assembled contig is long enough to cover multiple gaps. Therefore, the full range of a true alignment is hard to recover by global, end-to-end alignment approach. When we applied local alignment, one contig’s true alignment would be broken into locally aligned segments. The aligned segments should be consistent in both orientation and order, as well as allocated within a reasonable range. However, some other irrelevant linked read groups are unavoidably recruited in the pre-assembled read set and form irrelevant contigs with short alignment signals. Thus, we need to remove those mappings which are unlikely to be right. To define the longest, putative continuing, mapping range from the local alignment info, we trim the mapping result to form aligned segments of each contig. Starting from the longest aligned segment which has the best alignment score, we check two flankings of this segment and attach the neighbors to it if the neighbors are in the right order and with a small distance away from it. In this way, we expand the mapping range gradually; aligned segments which are located far from the main longest aligned segment will be considered as independent events and excluded. As a result, the remaining aligned segments would define a mapping range on a scaffold, which represent the contig’s expanded mapping segment.

##### III. Filling gaps in scaffolds

After the contig’s expanded mapping range is set, we check on every gap and see whether it can be filled or not. The workflow of gap filling is illustrated in Fig. 3. First, we classify gaps according to the distance between mapped contigs’ position and gaps’ position. For each aligned qualified contig (contig length >= 1k and coverage >= 2), we record the start and end positions of the longest contig mapping range on the scaffold. If a gap is located in between, we mark it as a fully covered gap. If one end of a gap is located outside the range but within a small distance (default of 50bp), we mark it as a partially covered gap. For those gaps which aren’t covered by any contig would be marked as unfilled and be skipped. The gaps can be marked as fully covered gaps and partially covered gaps at the same time due to the judgement based on different contigs’ mapping results. Contigs which fully cover given gaps for gap filling are considered to provide stronger evidence since they have two high quality alignments from both sides of the gap. Hence, the priority of a fully covered gap is higher and would be processed as the first-tier candidates. When there are multiple contigs fully covering the gap, the one with higher mapping score will be used first. In this case, we’re not able to remove the whole gap, nevertheless, the partially gap-covering contig will be able to increase scaffold content. As we are interested in the information which can be used to fill the gaps, the next step is to extract the unmapped part of contigs. For a fully covered gap, we find the nearest alignment of two flanks within a distance (default of 50bp) and check the mapping identity to make sure that contig maps properly to the scaffold. If a gap-covering contig does contain segment highly similar with mapping identity larger than 80 to both flankings of the gap, we believe that there are corresponding segments in that contig. Then, we find out that segments of contig and substitute for the gap. Similar to a fully covered gap, a partially covered gap has a flanking with high similarity (> 300 base match with mapping identity > 90) to part of a contig while the contig’s mapping range can’t cover the whole gap, so that we might only fill a part of gap by the segments which is unmapped are suitable to fill the gap. We further classify fully covered gaps into three groups (Supplementary Figure 6). “Perfect contigs’’ stand for the ideal group which contigs are normally mapped to a gap-containing region of a scaffold and the unmapped fractions corresponds to gaps; “Inconsistent contigs” means the unmapped fractions are not only found in the gap but also some extra segments (more than 50 bps) scattered in the contig that failed to align to the scaffold; “Conflict contigs” refers to the mapping problem that the two flankings of the gap are duplicated sequences which cause ambiguous conflicts in alignment or infers a collapse on repeats in the assembled contig. Inconsistent and conflict contigs are marked out instead of using them because we assume the genome assembly we have is mostly correct and would not prefer a substitution on the known sequence with newly assembled contigs. To avoid introducing the mis-assembled contig into the previous assembled genome, only the perfect contigs are capable of filling gaps. If a fully covered gap isn’t covered by any perfect contig, it can’t be fully filled and might be partially filled if there exists contig covering it partially. A pseudocode for the entire algorithm is provided in Supplementary Note 6.

#### 8.2 Global-Contig-Based (GCB) Gap Filling

This program can be viewed as an alternative for LCB Gap Filling. Instead of locally de novo assembling contigs, we fill the gaps via scaffolds using “G-contigs” as in Global-Contigs. G-contigs are existing sequences either from an earlier stage of the GABOLA pipeline or produced by other sequence assemblers. Long reads from Third-Generation Sequencing such as Pacbio SMRT Sequencing and Oxford Nanopore Technologies are also compatible as G-contigs (Supplementary Figure 5b).

During GCB Gap Filling, because G-contigs may be scaffolds containing a certain amount of gaps, we detect gaps that are longer than a specific length (default is 20) in G-contigs and break them into smaller segments to enhance the quality of the assembly. These segments are then used to fill in gapped regions, following the same process of point II and III from Section 8.1.

#### 8.3 Local-Contig-Based (LCB) Scaffolding

##### I. Assembling contigs

For each candidate scaffold pair, we created a barcode list containing their shared barcodes. Then, we collected all reads that have the same barcodes, whether they are mapped onto the scaffolds or not, to construct our L-contigs. We believe that we would gain more information with this approach, for short reads belonging to the same 10x GEM partition may be linked physically and provide us with long-range information of the original DNA fragment. Therefore, the more reads we recruited, the higher the quality of our assembled contigs might be. As for the assembling process, we use SPAdes assembler^82^ which is a genome assembly algorithm based on k-mer to build multi-sized de Bruijn graphs and construct contigs by graph-theoretical operations.

##### II. Concatenating scaffolds with high-quality contigs

With these L-contigs, we aligned them to the draft assembly with BWA-mem. We then checked the alignment within 20kbp of the associated end of each candidate scaffold pair. Within the 20kbp, we first kept those contigs with mapped length greater than 1 kbp and mapping identity greater than 70%. Subsequently, we began to find contigs that may truly connect the scaffold pair by locating the mapping position of the contigs. If the distance between the mapped position and the scaffold end is longer than one-fifth of the contig’s length, the contig is discarded. Finally, we concatenated the candidate scaffold pairs with the filtered contigs in the descending order of their shared barcode numbers (Supplementary Figure 5c). A pseudocode for this algorithm is provided in Supplementary Note 7.

#### 8.4 Global-Contig-Based (GCB) Scaffolding

This program can be viewed as an alternative for LCB Scaffolding. The process is fundamentally the same as point I and II stated in section 8.3 with a few alterations: Firstly, in place of L-contigs, we take G-contigs (see section 8.2) as input. Secondly, we changed the alignment tool from BWA-mem to minimap2 on account of its specialty in long read alignments (Supplementary Figure 5d).

### 9. Gene Annotation

Genome constructed by different genome assembly pipelines would be aligned to GRCh38 reference genome to identify increased genomic contents in GABOLA compared to Supernova assembly, and sequences in gap regions in reference genome. We applied minimap2 to perform alignment and further identified increased genome contents via direct genome comparison based on GRCh38 coordinate. The increased genome contents were then annotated based on GENCODE v29^25^.

### 10. Repeat Element Identification

To evaluate the performance of different assembly pipelines in the repeat regions, we applied RepeatMasker to identify the repeat elements and their corresponding repeat patterns. RepeatMasker was run with -species human setting. The outputs of RepeatMasker were further processed by perl script onecodetofindthemall.pl to categorize the repeat elements into several repeat patterns and generate copy number estimates^83^.

### 11. Gene Prediction for unmapped NS

In order to investigate whether the unmapped NS are genuinely exist in the genome, we applied Augustus^28^ to make gene prediction. We assume that if there are inferred genes located in these NS, these NS may possibly exist in the genome. Augustus was run with --*species*=*human*--*UTR*=*on setting*. To further validate these inferred genes, we performed protein BLAST^29^ to search if the predicted sequences are conserved in organisms.

