## Supplemental Information for "GABOLA: A Reliable Gap-Filling Strategy for *de novo* Chromosome-Level Assembly"

### Supplementary Information

1. **Supplementary Notes**
   Supplementary Note 1. Initial assembling of LLD0021C via Supernova
   Supplementary Note 2. Bionano Hybrid Scaffold on LLD0021C
   Supplementary Note 3. Initial assembling of CHM13 via Supernova
   Supplementary Note 4. Effect on gap filling regarding duplicate reads
   Supplementary Note 5. Testing out different barcode selection strategies
   Supplementary Note 6. Algorithm of LCB Gap Filling
   Supplementary Note 7. Algorithm of LCB Scaffolding
   Supplementary Note 8. Docker Manual
   Supplementary Note 9. Technical Notes

2. **Supplementary Figures**
   Supplementary Figure 1. Workflow for preprocessing FASTQs
   Supplementary Figure 2. Effects of marking duplicate reads
   Supplementary Figure 3. Gap filling result for 4 scaffolds under different BS affinity with duplicate and non-duplicate reads set
   Supplementary Figure 4. Three different barcode selection strategies
   Supplementary Figure 5. Illustration for the basic concept of the four main GABOLA modules
   Supplementary Figure 6. Workflow for filling gaps in scaffolds

3. **Supplementary Tables**
   Supplementary Table 1. Intra-scaffold gaps filled by GABOLA on LLD0021C
   Supplementary Table 2. Intra-scaffold gaps filled by GABOLA on CHM13
   Supplementary Table 3. Scaffold-based gap filling analysis of scaffold 21, 28, 36 and 39
   Supplementary Table 4. Analysis of barcoded reads against draft assembly after marking out duplicate and remove barcodes with under 3 read pairs
   Supplementary Table 5. Scaffold-based gap filling analysis of scaffold 21, 28, 36 and 39 using non-duplicate reads
   Supplementary Table 6. Gap filling analysis for each partition algorithm

### 1. Supplementary Notes

#### Supplementary Note 1. Initial assembling of LLD0021C via Supernova

```
> supernova run --id=LLD0021C_10x_supernova2 --
fastqs=10x_fastq/LLD0021C --localcores=64 --localmem=1024 &

> nohup supernova mkoutput --
asmdir=LLD0021C_10x_supernova2/outs/assembly --
outprefix=human_LLD0021C_10k_pseudohap --style=pseudohap --
minsize=10000 &
```

#### Supplementary Note 2. Bionano Hybrid Scaffold on LLD0021C

```
perl hybridScaffold.pl -n human_LLD0021C_10k_pseudohap.fasta \
-b LLD0021C.exp_refineFinal1_merged_q.cmap \
-c hybridScaffold_config.xml \
-r
/usr/local/bionano/Solve3.4_06042019a/RefAligner/8949.9232rel/RefAlig
ner \
-o IBMS_LLD0021Cpseudohap \
-f \
-B 2 \
-N 2
```

#### Supplementary Note 3. Initial assembling of CHM13 via Supernova

```
> supernova run --id=CHM13_supernova2 --fastqs=CHM13/10x_fastq --
localcores=64 --localmem=1024 &

> supernova mkoutput --asmdir=CHM13_supernova2/outs/assembly --
outprefix=CHM13_10k_pseudohap --style=pseudohap --minsize=10000 &
```

#### Supplementary Note 4. Effect on gap filling regarding duplicate reads

In order to find out relevant barcodes for each gap in each scaffold, we first mapped the trimmed read pairs and extracted barcode candidates with mappable reads on a given scaffold. Next, we recruited all read pairs in the barcode candidate list, regardless of whether they mapped to the target scaffold or not in the previous step, to form a read pair pool for gap filling. Due to the fact that repeats and duplicate sequences are present in a genome, we may be misled to recruit barcodes that are not related to a particular scaffold but have few reads mapped on it. Besides, a short genome fragment packed in a barcode GEM or not properly packed fragment would end up to low read pair throughput barcodes. These barcodes are unlikely to help us with assembling contigs well but will increase computation costs as well as noise info that leads to mis-assembling.

Thus, we try to set rules to exclude those troublesome barcodes. We start by defining a criterion for barcode quality: 1) a read is mapped if the overall mapping length is above 70% of its trimmed length, 2) a proper pair is a read pair with both reads being mapped on the same scaffold, 3) a unique (proper) pair, is a proper pair that map to one and only one scaffold. If a barcode relates to a scaffold, it must have unique reads on it. Here we introduce the number of unique pairs as a measurement for the affinity of each barcode-to-scaffold match, barcode-scaffold affinity (Briefly, BS affinity). Then we set different BS affinity to recruit different barcode candidates list/ read-pair pools, then we investigate the effects of BS affinity on gap-filling with scaffold-based partition algorithms. For example, barcode-scaffold affinity threshold setting to 5 means we want barcodes qualified to the unique-pair constraint by 5, i.e., at least 5 unique proper pairs to support the barcode-scaffold match. We select four scaffolds, scaffold 21, 28, 36, and 39 from our assembled eel genome draft as toy examples.

These scaffolds are 5.2, 4.8, 4.3 and 4.2 Mbps in length with 92.65%, 94.89%, 90.60% and 87.90% un-gapped base pairs respectively. For each gap, the most continuous aligned assembly from the relevant read pair pool is expected and applied to fill it out. When we set the BS affinity threshold higher, the number of both recruited barcodes and recruited read pairs decreased, and the computation time for the assembling/ gap filling task was shortened. As shown in Supplementary Table 3, BS affinity by 5 leads to better gap-filling results in scaffold 28 and 39 in terms of the percentage of filled gap length, while scaffold 21 and 36 get best performance in BS affinity by 1. The existence of gapped content affects the mappable read number of each barcode. BS affinity may relate to characteristics of gapped regions on each scaffold, and different constraints lead to varied results for it.

The outcomes of locality assembly are highly related to gap filling results. We found gap filling results vary greatly with different BS affinity. However, we also notice the variation may come from the artefact of duplicated read pairs. The read depth may be quite unevenly distributed along the scaffold (Supplementary Figure 2A). The proper pair number duplicated read pairs may be affected by duplicated read pairs (Supplementary Figure 2B). Thus, we tried to mark out duplicate read pairs in each barcode to give fair measurements of BS affinity. For any two read pairs from the same barcode, we mark the shorter or the latter read pairs as a duplicated read pair if their mapping positions are in the same start /end site and the sequence of reads are identical or prefixed identical to another read pair's sequence. Duplicated read pairs will be discarded so that every read pair from the same barcode is distinct. Indeed, we found that the read pair number of barcodes changes greatly in those high read number barcodes, and the accumulations of barcodes separated with proper pair threshold after mark duplicates procedure (Supplementary Figure 2C & 2D). The non-duplicated read set and gap filling results using non-duplicate barcoded read pairs are shown in Supplementary Table 4 & 5. After collapsing duplicated read pairs, recruited barcodes and read pairs are all slightly reduced, and the computation time also reduces slightly in most of the test combinations. The best performance of gap filling is shifted toward the unique-pair constraint of 1 or 3. The percentage of filled gap length increased in scaffold 21 (+6.7%), scaffold 28 (+8.6%), and scaffold 39 (+3.4%) except a small decrease in scaffold 36 (1 % decreased). Although the number of recruited barcodes and read pairs are smaller, the difference in read-pair constraints is diminished and the performance of our gap filling approach in scaffold 21 improved. Supplementary Figure 3 shows the performance of this four-scaffold gap filling test. We found excluding duplicates does improve the data quality that helps fill more gaps and reach better results.

**Supplementary Note 5. Testing out different barcode selection strategies**

We apply this model to the whole eel genome assembly (eel v4.0, n=948 to catch 95% of expected genome size, length sum: 991.45M, with unfilled gap #N: 4,372,491) to see the improvement of genome assembly. For recruiting barcodes with strong affinity to the target scaffold, we first set a BS affinity in 3 to create barcode lists for each scaffold first. All trimmed reads from the barcode list of a given scaffold are used to be assembled and then to fill the gaps. We found the enormous number of recruited reads result in exhaustive runtime. The longest scaffold is about 13.5Mbp, and it costs more than ten days for assembling. We aborted this gap filling task and consider this scaffold-based partition would be an inefficient choice for longer scaffolds. To solve this problem, algorithms adopting two new partition strategies, gap-based partition and window-based partition, are designed (Supplementary Figure 4). For a gap-based partition, we extracted a sub list from the barcode list for each gap based on the gap's position. Gap-related barcode sub lists are defined by barcode-scaffold affinity score but we modified the BS affinity score to gap's flanking region only. For window-based partitions, gaps located within a window size (60,000bp) are set as a group and the extended flanking boundary (5,000bp) of this group would not contain any gap. We move back to the four-scaffold example set to test our approach. Supplementary Table 6 shows the comparison of the gap filling result in the three partition strategies. All benchmarking was done on a 96 CPUs and 1.48T RAM machine. Obviously, gap-based and window-based partitions decrease input size and implementation of parallelizing jobs would shorten the runtime for each job. Percentage of filled gap length of the three partition algorithms is gap-based > window-based > scaffold-based, suggesting that strategies to recruit more localized barcoded reads also lead to better gap-filling results.

#### Supplementary Note 6. Algorithm of LCB Gap Filling

```
STEP I. Barcode Selection
C70M60 sam file: The alignment of 10x linked reads to our draft assembly with mapping
identity greater than or equal to 70% and mapping quality greater than or equal to 60.
{BXlist}S: Barcode list for each scaffold.
{BXlist}S.G: Barcode list for each gap per scaffold.

PROGRAM ProduceBXList (C70M60 sam file, draft assembly):
 Rename scaffolds by descending order of size;
 FOR each scaffold DO
  IF (number of mapped read pairs >= 3)THEN
   Add barcode to {BXlist}S;
  ENDIF
 ENDFOR
 FOR each gap on scaffold DO
  Determine flanking region based on gap size;
  IF (number of mapped read pairs on gap's flanking >= 2)THEN
     Select barcode from {BXlist}S and add barcode to {BXlist}S.G;
   ENDIF
 ENDFOR
 IF (number of barcodes in {BXlist}S.G <= 10)THEN
   Gap will not be processed;
 ENDIF
END
```

```
STEP II. De novo Local Assembly of Contigs
Nondup fastq: Fastq file of non-duplicated 10x reads split according to barcode type.

PROGRAM Assemble (nondup fastq):
 Determine number of gaps to process simultaneously;
 FOR each gap
  DO Collect barcoded reads based on {BXlist}S.G from nondup fastq;
       Assemble reads into contigs;
 ENDFOR
END
```

**STEP III. Gap Filling**

*Lc*: Left-most position of the longest contig mapping range on a scaffold.
*Rc*: Right-most position of the longest contig mapping range on a scaffold.
*I*: Set of longest contig mapping intervals.
*MI*: Mapping identity of contig on scaffold.
*ML*: Length of matched base of contig on scaffold.
*U*: Set of unmapped segments of contig on scaffold.

```
PROGRAM Fill (draft assembly):
 FOR each contig DO
  IF (contig length >= 1kbp  contig coverage >= 2)THEN
     Align contig to scaffold;
     IF (contig is not within the flankings of any gap) THEN
     Remove contig;
   ELSE
     {Lc, Rc} = longest mapping range of contig on the scaffold;
     I = I  {Lc, Rc};
   ENDIF
  ENDIF
 ENDFOR
 FOR each gap DO
   IF (gap  {Lc, Rc}) THEN
     Gap is considered fully-covered;
     FOR each contig covering the gap DO
        WHILE ( MI >=80 on both flankings of gap) DO
         IF ( U is only within the gap) THEN
          Gap is fully filled;
         ELSE
          Gap will not be filled;
         ENDIF
        ENDWHILE
    ENDFOR
   ELIF (gap  {Lc-50bp ,Rc}  {Lc, Rc+50bp}) THEN
     Gap is considered partially-covered;
     FOR each contig partially-covering the gap DO
        WHILE ( MI >=90 on the flanking of gap  ML >=300bp ) DO
         Fill part of the gap;
       ENDWHILE
     ENDFOR
   ELSE
     Gap is considered unfillable;
   ENDIF
 ENDFOR
END
```

#### Supplementary Note 7. Algorithm of LCB Scaffolding

**STEP I. Determine candidate scaffold pairs**

*E*: Candidate of scaffold end pairs.
*L*: Barcode list on each scaffold head and tail.
*F*: Filtered candidate of scaffold end pairs.

```
PROGRAM Preprocessing (E,L):
 FOR each scaffold pair in E DO
     ScafA, ScafB = scaffold end pair
        IF (ScafA==ScafB) THEN
          Remove scaffold pair;
        ELIF (ScafA or ScafB appear more than two times in E) THEN
          Add only the scaffold pairs with first and second most
          barcode counts to F ;
        ENDIF
 ENDFOR
 FOR each scaffold pair in F DO
     ScafAf,ScafBf = scaffold end pair;
     Collect barcode list on ScafAf end from L;
     Collect barcode list on ScafBf end from L;
  ENDFOR
END
```

**STEP II. *De novo* Assembling of Contigs**

*F*: Filtered candidate of scaffold end pairs.
Nondup fastq: Fastq file of non-duplicated 10x reads splited according to barcode type.

```
PROGRAM Assemble(F, nondup fastq):
   Determine number of scaffold pairs to process simultaneously;
   FOR each scaffold pair in F DO
      ScafAf,ScafBf = scaffold end pair;
      Collect reads from nondup fastq based on barcode list on ScafAf;
      Collect reads from nondup fastq based on barcode list on ScafBf;
      Assemble reads into contigs;
   ENDFOR
END
```

**STEP III. Scaffolding**

*F*: Filtered candidate of scaffold end pairs.
Cf: Candidate contigs for each scaffold pair in *F*
*MI*: Mapping identity of contig on scaffold.
*ML*: Length of matched base of contig on scaffold.

```
PROGRAM InterScaffolding (F, draft assembly):
   FOR each scaffold pair in F DO
      Align Cf to draft assembly;
      FOR each contig in Cf DO
         WHILE (contig is aligned within 20kbp of both scaffold ends)
         DO
            IF ( ML >= 1kbp and MI >= 70) DO
               Concatenate two scaffolds;
            ENDIF
         ENDWHILE
      ENDFOR
 ENDFOR
END
```

#### 2. Supplementary Figures

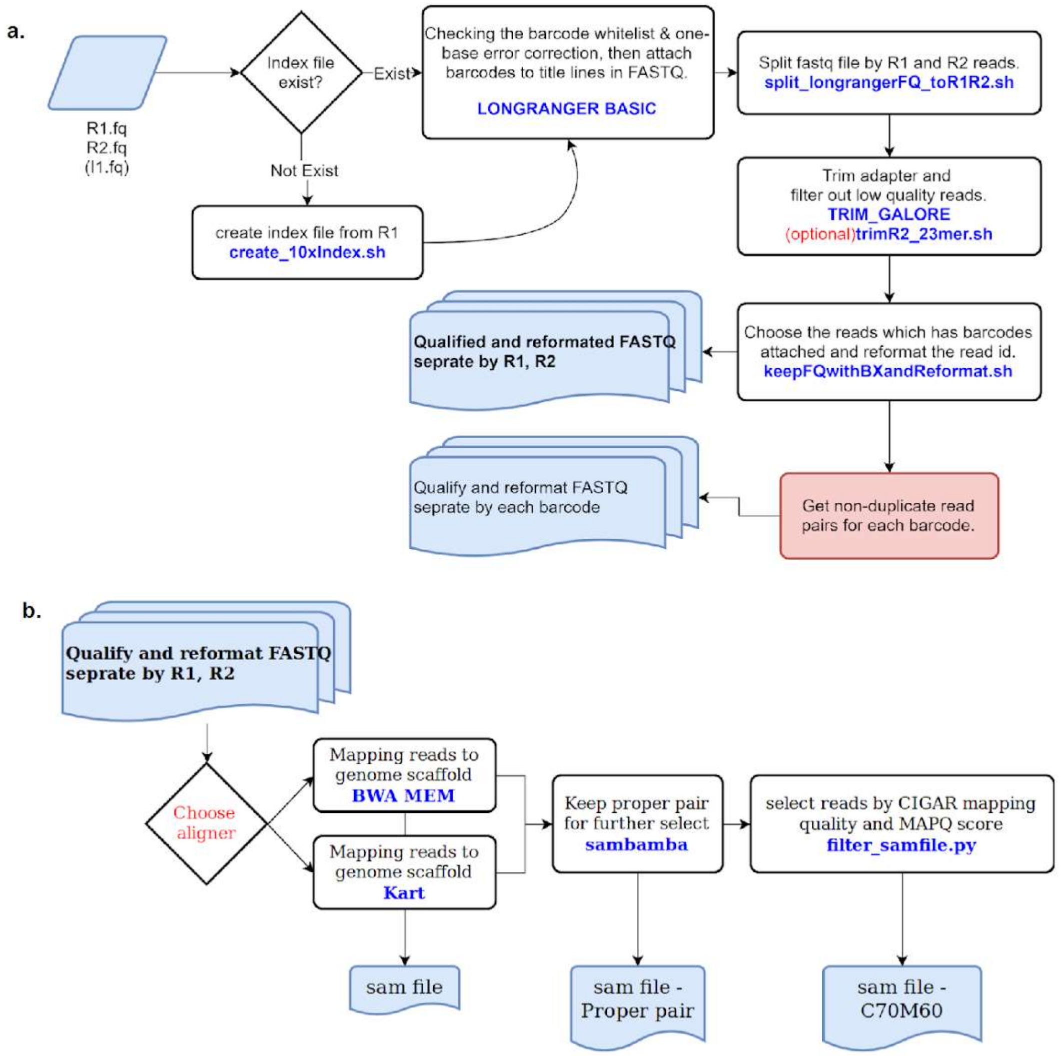


**Supplementary Figure 1. Workflow for preprocessing FASTQs**

**a.** We process the raw linked reads by taking FASTQ files created by longranger mkfastq and perform barcode processing including error correction, barcode white-listing, and attaching barcodes to reads. Then, we use TrimGalore to apply adapter and quality trimming to fastq files. After that, you can choose to trim the 23-mer of R2 prefix or not. Finally, we process all fastq files to remove duplicate reads.

**b.** We map reads to the genome and filter them: We begin with aligning reads to genome scaffolds by BWA MEM or Kart. Then, we keep proper pairs by SAMBAMBA (v0.6.9). Finally, we filter the alignment by match quality calculated from CIGAR and MAPQ score. #definition of proper pair: mapped and not (secondary alignment or duplicate or supplementary or chimeric)

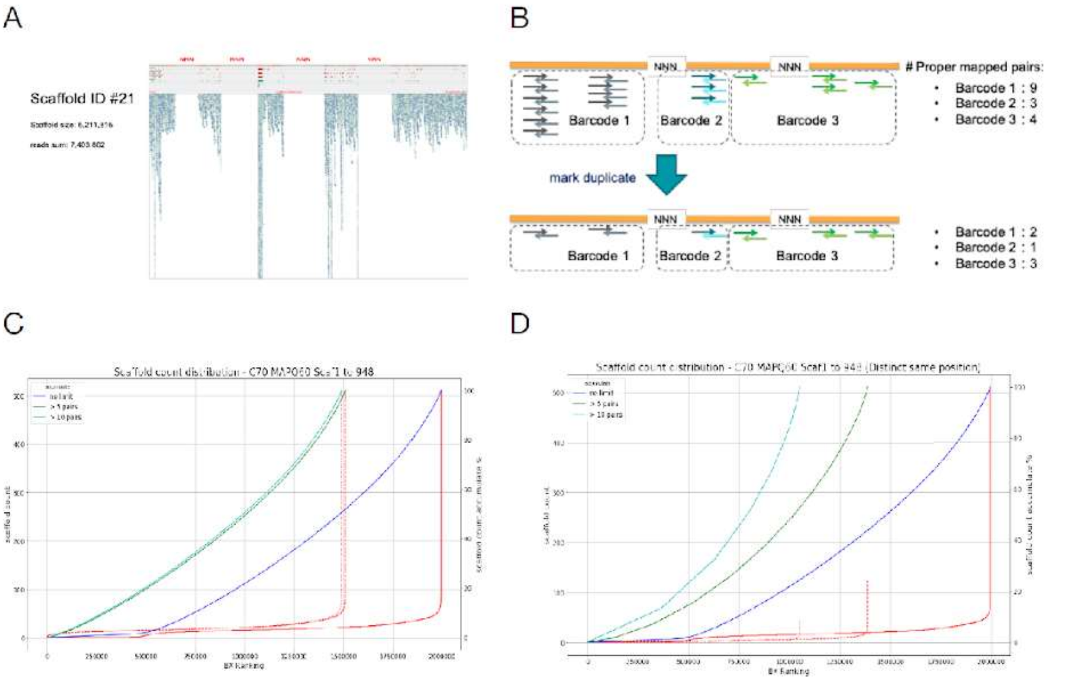


**Supplementary Figure 2. Effects of marking duplicate reads**

**A.** Visualizing the reads mapped on the first 25k region of scaffold 21 by IGV. The read depth is quite unevenly distributed along the scaffold.

**B.** A pilot example of mark duplicate. Read pairs from a barcode were counted once if they are located in the same start/ end base position of a scaffold and in the same direction. Mark duplicate would work like a collapse of redundant read pairs and give a fair estimation of barcode-to-scaffold affinity.

**C.** Barcode-to-Scaffold Assignment from the mapping result of original 10x linked reads. Barcodes are sorted ascendingly by the number of mappable scaffolds without limitation or set threshold to 5 or 10 high-quality proper pairs. Line in red is the scaffold number of each barcode and blue/green/cyan lines present the accumulation of barcodes in percentage.

**D.** Barcode-to-Scaffold Assignment from the mapping result of mark-duplicated 10x linked reads. All are similar to panel C except the data is derived with the read pair collapsing for unique location of read pairs.

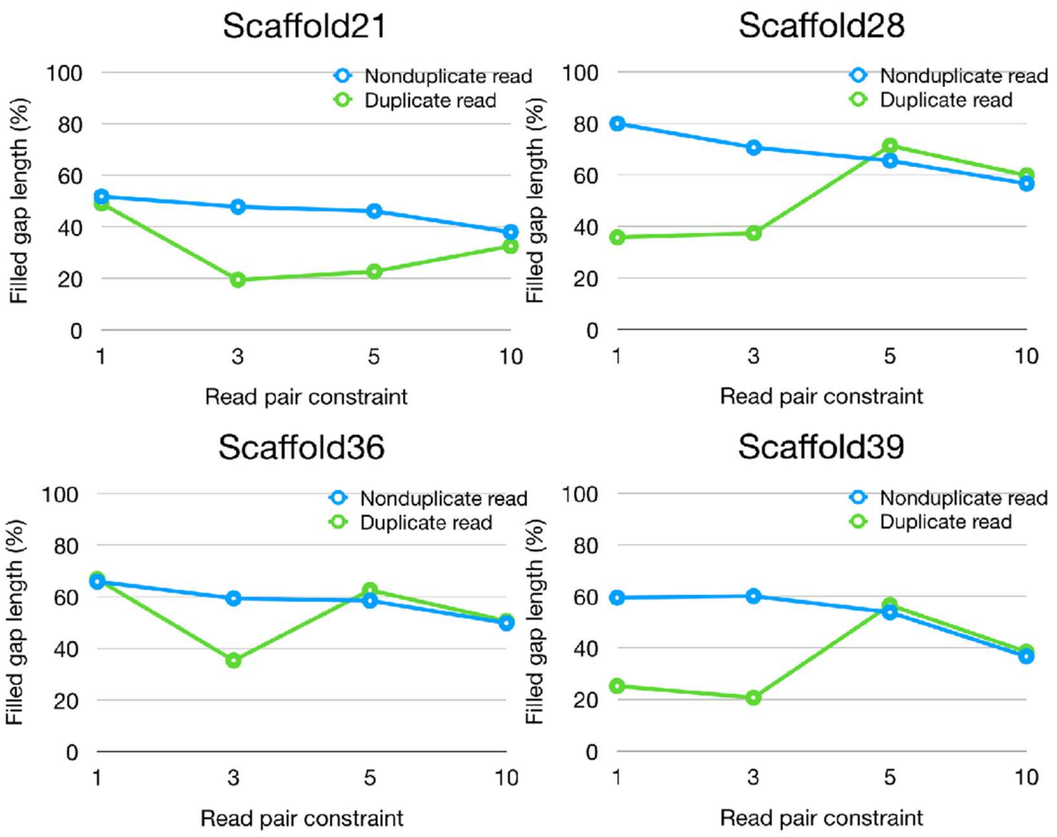


**Supplementary Figure 3. Gap filling result for 4 scaffolds under different BS affinity with duplicate and non-duplicate read set.**

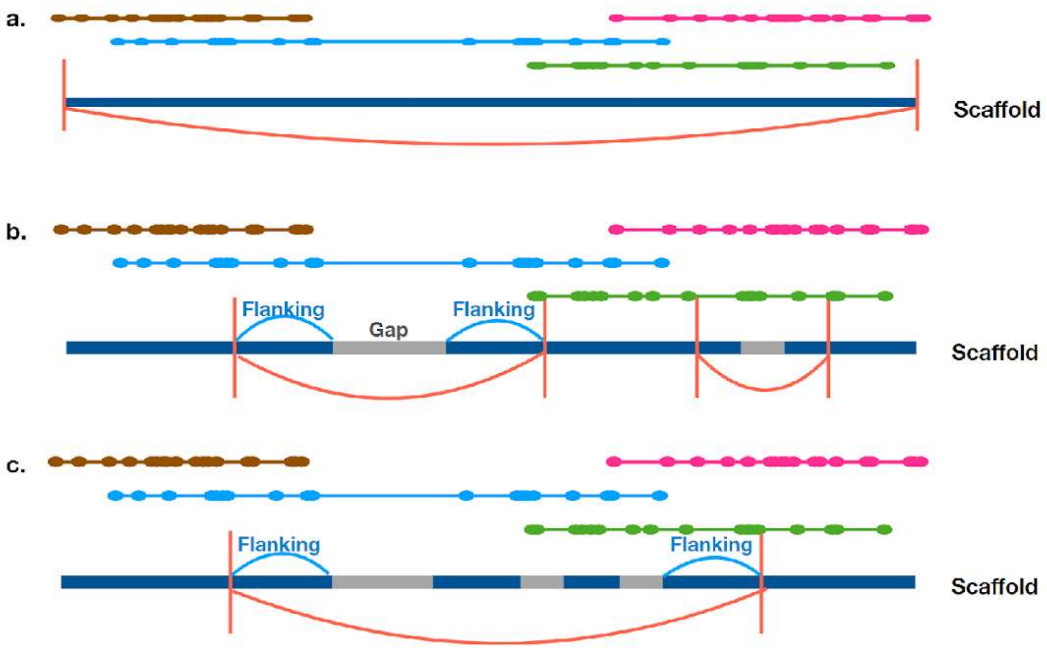


**Supplementary Figure 4. Three different barcode selection strategies**

**a.** Scaffold-based barcode selection: We collect barcodes mapped onto the entire scaffold

**b.** Gap-based barcode selection: We collect barcodes mapped onto the flanking region of each gap. The flanking size is determined by the gap size.

**c.** Window-based barcode selection: We group gaps located within a window of 60kbp and collect barcodes mapped onto the flanking region (5 kbp) per window.

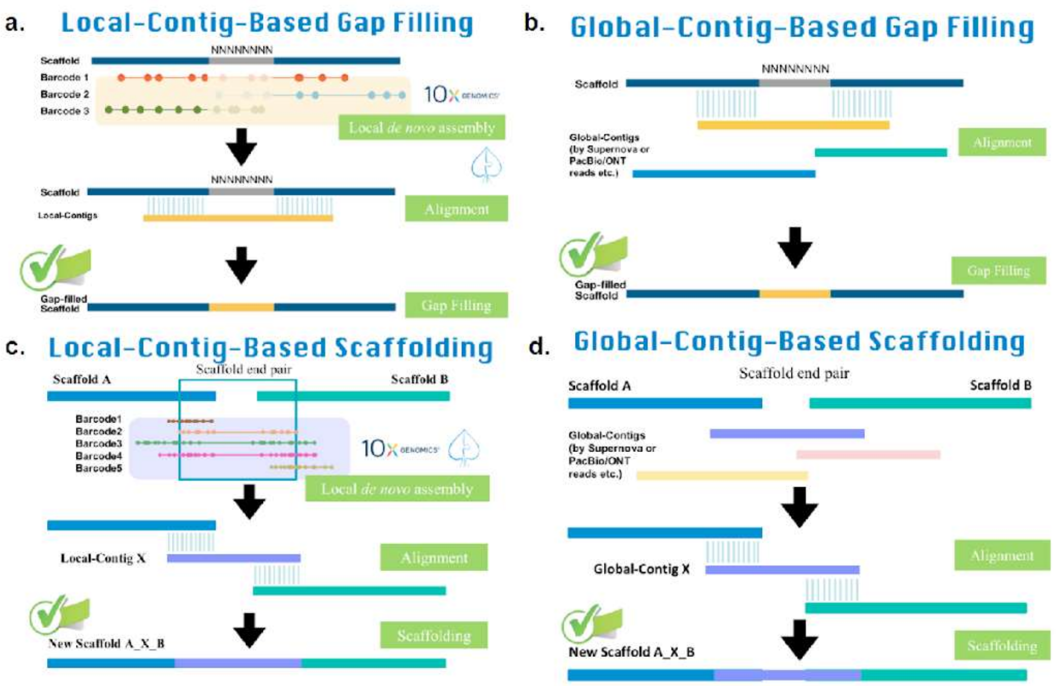


**Supplementary Figure 5. Illustration for the basic concept of the four main GABOLA modules**

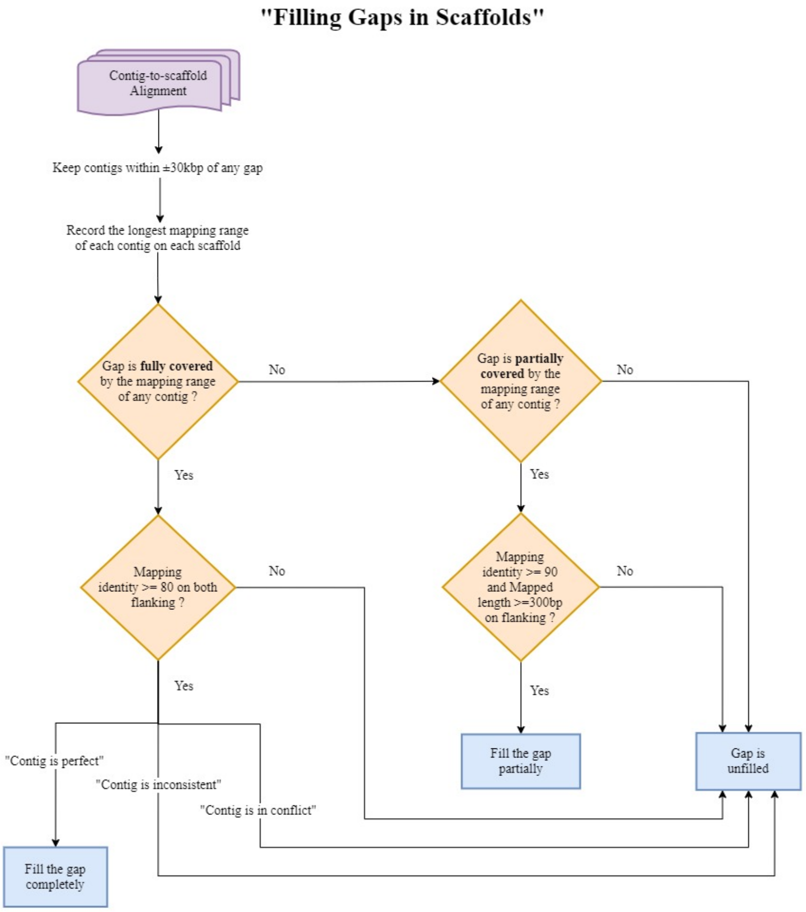


**Supplementary Figure 6. Workflow for filling gaps in scaffolds**

First, gaps are classified into three types based on contig-gap flankings alignments. For those fully-covered gaps, we check the mapping quality of the extended contig segment to the gap flankings, and see whether it meets the criteria in both flanking to replace the gap regions. To avoid introducing the misassembled contig into the previous assembled genome, only the perfect contigs are capable of filling gaps.

#### 3. Supplementary Tables

**Supplementary Table 3. Scaffold-based gap filling analysis of scaffold 21, 28, 36 and 39**

| | BS affinity | #Recruited Barcode | #Recruited Read Pairs | Wall Clock Time (hr) | Filled Gap Length (%) |
|---|---|---|---|---|---|
| **Scaffold21 {dup}**<br>• **5.2Mbps,**<br>• # Gaps: 313<br>• **# N: 383.3Kbps** | 1 | 178,719 | 69,002,194 | 70.6 | **45.1** |
| | 3 | 86,708 | 35,721,283 | 40.7 | 19.5 |
| | 5 | 53,314 | 20,582,022 | 24.1 | 22.7 |
| | 10 | 20,176 | 8,997,030 | 8.4 | 32.6 |
| **Scaffold28**<br>• **4.8Mbps,**<br>• **# Gaps: 244**<br>• **# N: 243.5Kbps** | 1 | 168,527 | 64,705,613 | 76.1 | 35.8 |
| | 3 | 86,588 | 35,488,861 | 36.5 | 37.4 |
| | 5 | 54,845 | 23,042,400 | 18.5 | **71.3** |
| | 10 | 22,122 | 9,720,551 | 9.4 | 59.8 |
| **Scaffold36**<br>• **4.3Mbps,**<br>• **# Gaps: 332**<br>• **# N: 402.3Kbps** | 1 | 146,548 | 56,652,569 | 50.6 | **66.9** |
| | 3 | 72,373 | 29,750,862 | 31.8 | 35.3 |
| | 5 | 44,868 | 18,949,958 | 17.9 | 62.6 |
| | 10 | 17,153 | 7,577,638 | 7.6 | 50.6 |
| **Scaffold39**<br>• **4.2Mbps,**<br>• # Gaps: 387<br>• **# N: 505.1Kbps** | 1 | 134,141 | 52,281,320 | 35.4 | 25.2 |
| | 3 | 62,129 | 25,699,474 | 28.9 | 20.7 |
| | 5 | 36,790 | 15,609,672 | 13.4 | **56.6** |
| | 10 | 13,024 | 5,735,802 | 4.7 | 38.5 |

**Supplementary Table 4. Analysis of barcoded reads against draft assembly after marking out duplicate and remove barcodes with under 3 read pairs**

| | # of Reads | # of Barcodes | #Scaffold with at least one mapped read |
|---|---|---|---|
| Barcoded read | 1,177,151,368 | 3,449,903 | - |
| Trimmed read | 1,142,645,930 | 3,423,181 | - |
| Non-duplicate read | 931,300,700 | 2,395,977 | - |
| Aligned read | 931,300,700 (100%) | 2,395,977 (100%) | 7,804 (100%) |
| Proper pair alignment | 652,825,738 (70.1%) | 2,262,747 (94.4%) | 7,801 (99.9%) |
| Mapping identity > 70 and Mapping quality == 60 | 280,540,246 (30.1%) | 1,772,060 (74.0%) | 6,126 (78.5%) |

**Supplementary Table 5. Scaffold-based gap filling analysis of scaffold 21, 28, 36 and 39 using non-duplicate reads**

| | BS affinity | #Recruited Barcode | #Recruited Read Pairs | Wall Clock Time (hr) | Filled Gap Length (%) |
|---|---|---|---|---|---|
| **Scaffold21**<br>● **5.2Mbps,**<br>● # Gaps: 313<br>● **# N: 383.3Kbps** | 1 | 177,302 | 64,370,474 | 74.1 | **51.8** |
| | 3 | 86,413 | 33,744,148 | 31.5 | 47.8 |
| | 5 | 53,087 | 20,893,778 | 22.3 | 46.1 |
| | 10 | 20,018 | 8,259,223 | 7.0 | 38.0 |
| **Scaffold28**<br>● **4.8Mbps,**<br>● **# Gaps: 244**<br>● **# N: 243.5Kbps** | 1 | 167,131 | 60,358,361 | 62.3 | **79.9** |
| | 3 | 86,336 | 32,925,945 | 31.6 | 70.6 |
| | 5 | 54,597 | 21,313,213 | 20.9 | 65.5 |
| | 10 | 21,965 | 8,929,363 | 9.6 | 56.6 |
| **Scaffold36**<br>● **4.3Mbps,**<br>● **# Gaps: 332**<br>● **# N: 402.3Kbps** | 1 | 145,305 | 52,877,734 | 48.1 | **65.9** |
| | 3 | 72,176 | 27,639,712 | 26.7 | 59.4 |
| | 5 | 44,677 | 17,541,650 | 16.5 | 58.5 |
| | 10 | 17,018 | 6,955,085 | 8.4 | 49.8 |
| **Scaffold39**<br>● **4.2Mbps,**<br>● # Gaps: 387<br>● **# N: 505.1Kbps** | 1 | 133,138 | 48,774,109 | 34.0 | 59.5 |
| | 3 | 61,929 | 23,840,394 | 21.1 | **60.1** |
| | 5 | 36,602 | 14,412,116 | 11.0 | 53.8 |
| | 10 | 12,914 | 5,257,810 | 5.6 | 36.7 |

#### Supplementary Table 6. Gap filling analysis for each partition algorithm

| **Partition** | Scaffold-based | | | | **Gap-based** | | | | Window-based | | | |
|---|---|---|---|---|---|---|---|---|---|---|---|---|
| **ScaffoldID** | Scaf21 | Scaf28 | Scaf36 | Scaf39 | Scaf21 | Scaf28 | Scaf36 | Scaf39 | Scaf21 | Scaf28 | Scaf36 | Scaf39 |
| **#Task** | 1 | 1 | 1 | 1 | 311 | 243 | 332 | 386 | 69 | 58 | 55 | 55 |
| **#Total Processed Barcode** | 86,707 | 86,588 | 72,373 | 62,128 | 82,837 | 66,143 | 89,343 | 86,289 | 70,280 | 62,097 | 63,542 | 54,707 |
| **#Total Processed Read Pair** | 35,721,245 | 35,488,861 | 29,750,862 | 25,699,346 | 35,181,566 | 27,996,758 | 37,579,213 | 36,519,860 | 29,524,954 | 25,985,365 | 26,489,125 | 22,908,558 |
| **Wall Clock Time (hr)** | 45 | 37 | 32 | 18 | 5.9 | 6.2 | 4.1 | 6.3 | 2.3 | 2.0 | 2.1 | 2.0 |
| **Filled Gap Length (%)** | 19.5 | 37.4 | 35.3 | 20.7 | **65.0** | **80.2** | **76.8** | **68.5** | 57.4 | 68.8 | 71.6 | 60.4 |
